# mRNA delivery of mosaic-8 pan-sarbecovirus RBD vaccines elicits distinct antibody epitope signatures

**DOI:** 10.1101/2025.10.21.683747

**Authors:** Alexander A. Cohen, Jennifer R. Keeffe, Lusineh Manasyan, Indeever Madireddy, Ange-Célia I. Priso Fils, Kim-Marie A. Dam, Haley E. Stober, Rory A. Hills, Woohyun J. Moon, Paulo J.C. Lin, Mark R. Howarth, Magnus A.G. Hoffmann, Pamela J. Bjorkman

## Abstract

Effective pan-sarbecovirus vaccines could prevent future zoonotic spillovers of SARS-like betacoronaviruses. We previously developed protein-based mosaic-8 nanoparticles displaying eight diverse sarbecovirus RBDs, either individually (mosaic-8 RBD-NPs) or as two “quartets” of four tandemly-arranged RBDs (dual-quartet RBD-NPs), which elicited broadly cross-reactive antibodies but require multi-component manufacturing. Here, we address scalability challenges by extending the mosaic-8 concept to mRNA by encoding membrane-bound RBD quartets as dual-quartet RBD-mRNA and dual-quartet RBD-EABR-mRNA, the latter leveraging ESCRT- and ALIX-binding region (EABR) technology for immunogen display on cell surfaces and secreted virus-like particles. Compared with protein-based mosaic-8 immunogens, mRNA-encoded mosaic-8 vaccines induced equivalent or enhanced antibody breadth, neutralization potencies, T-cell responses, and targeting of conserved RBD epitopes. In addition, mRNA-encoded mosaic-8 vaccines elicited more balanced IgG subclass profiles and increased Fcγ receptor–binding IgGs, consistent with potentially superior Fc effector functions. These findings demonstrate successful translation of mosaic-8 RBD-NPs into mRNA/EABR-mRNA vaccines, enabling scalable manufacturing and improving protection against future sarbecovirus outbreaks. Finally, our newly developed technique, Systems Serology–Polyclonal Epitope Mapping (SySPEM), revealed distinct IgG-subclass-specific epitope signatures across mRNA, EABR-mRNA, and protein vaccines, demonstrating that the mode of antigen display can shape epitope recognition.

**Summary:** We translated a pan-sarbecovirus RBD vaccine from protein nanoparticles to scalable mRNA and EABR-mRNA platforms encoding RBD quartets. Compared with protein-based immunogens, mRNA-based vaccines matched or improved antibody breadth, T-cell responses, Fc functionality, and conserved epitope targeting. A newly-developed Systems Serology–Polyclonal Epitope Mapping (SySPEM) technique revealed that antigen presentation modality shapes IgG subclass–specific epitope recognition.

## Introduction

Three coronaviruses spilled over from animal hosts to cause human epidemics or pandemics in the past 25 years: SARS-CoV (hereafter SARS-1), MERS-CoV, and SARS-CoV-2 (hereafter SARS-2) (Andersen et al., 2020). Two of these, SARS-1 and SARS-2, are members of the SARS-like betacoronavirus (sarbecovirus) subgenus. SARS-2 continues to infect humans, driving emergence of new variants that necessitate regular updates of current COVID-19 vaccines (Roederer et al., 2024). Since we cannot predict how SARS-2 will continue to mutate or which zoonotic sarbecovirus will next infect humans, we need a pan-sarbecovirus vaccine that does not require updating to provide broad protection against new SARS-2 variants and future sarbecovirus spillovers (Whittaker et al., 2025).

Spike trimers of coronaviruses function in host cell entry after one or more receptor-binding domains (RBDs) adopt an “up” position to permit interactions with a host receptor (Hills et al., 2025). RBD targeting has been suggested for COVID-19 vaccine development as RBDs are the main target of neutralizing antibodies (Abs) (Kleanthous et al., 2021). We previously described a vaccine approach that elicits broad Ab responses to diverse sarbecoviruses involving RBDs from multiple sarbecoviruses that are covalently linked using the SpyCatcher-SpyTag system (Bruun et al., 2018) to a SpyCatcher-mi3 protein nanoparticle (NP) (Cohen et al., 2021a; Cohen et al., 2024; Cohen et al., 2022; Wang et al., 2025). In this approach, mosaic-8 RBD-NPs displayed eight different sarbecovirus spike RBDs attached randomly to the 60 positions of the mi3 NP (Fig. 1, A and B). We hypothesized that B cells bearing cross-reactive receptors (B cell receptors; BCRs) recognizing features in common between adjacent non-identical RBDs are preferentially activated compared to B cells with BCRs recognizing immunodominant strain-specific epitopes (Cohen et al., 2022) (Fig. 1 A). To investigate this, we used deep mutational scanning (DMS) (Starr et al., 2020) to map RBD epitopes recognized by Abs from animals immunized with mosaic-8 RBD-NPs or homotypic RBD-NPs (only presenting SARS-2 RBD), revealing targeting of more conserved RBD regions by mosaic-8 RBD-NP–elicited Abs compared with targeting of more variable RBD regions by homotypic RBD-NP–elicited Abs (Cohen et al., 2025; Cohen et al., 2024; Cohen et al., 2022; Hills et al., 2024; Wang et al., 2025). This finding rationalized results from challenge experiments in which mosaic-8 RBD-NPs protected from both a matched viral challenge (from a virus represented by an RBD on the NP) and a mismatched challenge (from a virus not represented by an RBD on the NP), whereas homotypic RBD-NPs protected only from a matched challenge (Cohen et al., 2022). We also found broader Ab responses elicited by mosaic-8 RBD-NPs in animals that were pre-vaccinated with COVID-19 vaccines (Cohen et al., 2024). These results suggest that a mosaic-8 RBD-based vaccine given as a COVID-19 booster could provide increased protection from SARS-2 variants and prevent future sarbecovirus spillover(s) from causing an epidemic or pandemic.

**Figure 1.**
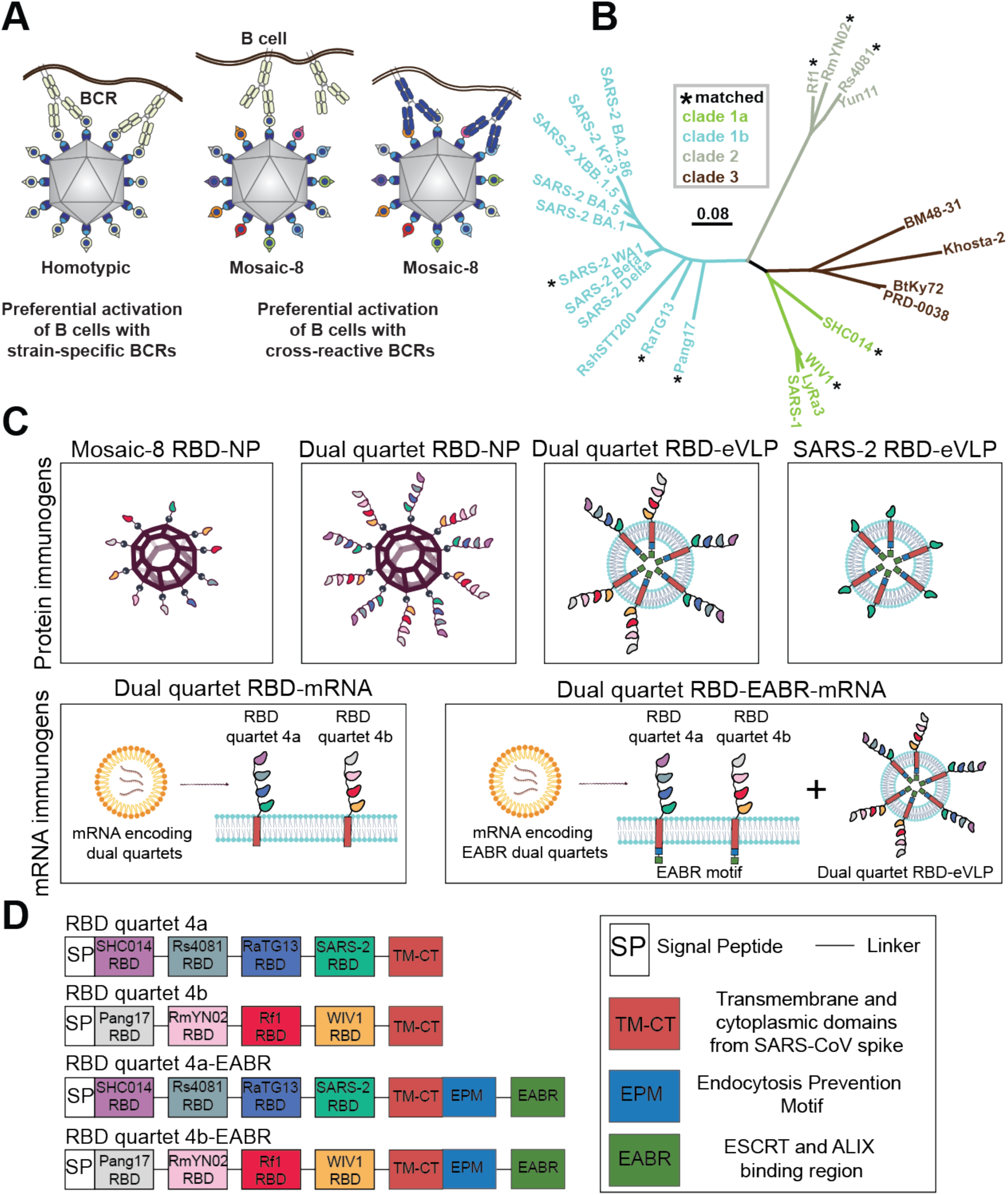
Protein- and mRNA-based pan-sarbecovirus immunogens. (A) Avidity hypothesis. Left: Membrane-bound strain-specific BCRs (pale yellow) using avidity to recognize a strain-specific epitope (pale yellow triangle) on antigens attached to a homotypic NP. Middle: Strain-specific BCRs binding with only a single Fab (i.e., not using avidity) to a strain-specific epitope (triangle) on an antigen attached to a mosaic NP. Right: Cross-reactive BCRs using avidity to recognize a common epitope (blue circle) presented on different antigens attached to a mosaic NP, but not to strain-specific epitopes (triangles). Only a fraction of the 60 attached RBDs are shown for clarity. (B) Sarbecovirus phylogenetic tree (made using a Jukes-Cantor generic distance model with Geneious Prime® 2023.1.2) calculated from amino acid sequences of RBDs aligned using Clustal Omega (Sievers et al., 2011). Viruses with RBDs included in RBD quartets and mosaic-8 RBD-NPs are indicated with an asterisk. Scale bar = phylogenetic distance of 0.08 nucleotide substitutions per site. (C) Schematics of protein-based and mRNA-based pan-sarbecovirus immunogens (not to scale; showing only a fraction of antigens for clarity). (D) Schematics of mRNA constructs encoding RBD dual quartet immunogens used in this study.

Manufacturing a mosaic-8 RBD-NP vaccine would require producing nine proteins: eight RBDs and a NP to which the RBDs are coupled. Making an mRNA-based version of a mosaic-8 vaccine would also pose challenges due to the requirement of synthesizing eight individual modified mRNAs. The RBD-encoding mRNAs could either be packaged into individual lipid nanoparticles (LNPs) or co-formulated into a single LNP. In the first case, some cells would not take up all eight mRNA LNPs, therefore presenting fewer instances of different adjacent RBDs, potentially reducing avidity effects to activate B cells with maximally cross-reactive BCRs (Fig. 1 A). In the second case, co-formulation of eight mRNAs into a single LNP would present technical and regulatory challenges. A potential solution relevant to converting a mosaic-8 vaccine into an mRNA format involves making mi3 NPs presenting two RBD “quartets” (each containing four RBDs arranged in tandem), which elicit equally broad and potent Abs as the original mosaic-8 RBD-NP presenting monomeric RBDs (Hills et al., 2024) (Fig. 1 C; dual quartet RBD-NP). Dual quartet RBD-NPs require manufacture of only three components (two quartets and a NP), and by synthesizing mRNAs encoding membrane-bound RBD quartets, could be more easily converted to an mRNA-LNP format than mosaic-8 RBD-NPs. In addition to simpler manufacturing, mRNA-LNP vaccines offer advantages over protein-based vaccines in that viral antigens undergo translation in the cytoplasm, thereby providing a source of viral peptides for presentation by MHC class I molecules to activate CD8+ cytotoxic T cells (Gote et al., 2023).

Conversion of dual RBD quartet immunogens to an mRNA format (Fig. 1 C; dual quartet RBD-mRNA) is also compatible with using the ESCRT- and ALIX-binding region (EABR) mRNA vaccine platform, which presents immunogens on cell surfaces and on circulating enveloped virus-like particles (eVLPs), thereby combining attributes of both mRNA and protein-NP vaccines (Hoffmann et al., 2023) (Fig. 1 C; dual quartet RBD-EABR-mRNA). eVLP formation is achieved by appending an EABR sequence to the cytoplasmic domain of the mRNA-encoded immunogen, which recruits cellular ESCRT proteins to induce eVLP budding from the plasma membrane. We previously showed that two immunizations with mRNA-LNPs encoding spike-EABR elicited potent CD8+ T-cell responses and superior neutralizing Ab responses against original and variant SARS-2 compared to conventional spike-encoding mRNA-LNPs, improving neutralizing titers >10-fold against Omicron-based variants (Hoffmann et al., 2023).

Until this study, it was not known whether presentation of membrane-bound RBD quartets on eVLPs and/or the surfaces of cells would elicit cross-reactive Abs to the extent we observed using RBD quartets or single RBDs covalently linked to rigid protein NPs. Here, we show that the advantages of protein-based mosaic-8 RBD-NPs can be transferred to an mRNA-LNP format. We compared immune responses in mice immunized with dual quartet RBD-mRNA versions of mosaic-8 immunogens (dual quartet RBD-mRNA and dual quartet RBD-EABR-mRNA) with responses to two protein-based mosaic-8 NPs (dual quartet RBD-NPs (Hills et al., 2024) and EABR-based dual quartet RBD-eVLPs) or to homotypic SARS-2 RBD-eVLPs. Immunization with mRNA immunogens elicited equivalent or improved binding breadths, neutralization potencies, T cell responses, and targeting of conserved RBD epitopes across sarbecoviruses compared to protein-based mosaic-8 immunogens, demonstrating successful conversion of the mosaic-8 RBD vaccine to mRNA formats that enable streamlined, scalable manufacturing of a pan-sarbecovirus vaccine.

## Results

### mRNA-encoded RBD quartets are presented on cell surfaces and eVLPs

mRNA-LNPs direct cell surface expression of a variety of gene products, including viral antigens (Hou et al., 2021). We investigated whether protein-based soluble dual quartets could be modified to induce cell surface expression by creating constructs that encode an RBD quartet followed by a linker, transmembrane anchoring sequence, and cytoplasmic tail (Fig. 1 D; RBD quartets 4a and 4b). The eight RBDs in the 4a and 4b quartet sequences correspond to the RBDs used in previous studies of protein-based dual quartet RBD-NPs (Hills et al., 2024) and to RBDs in a mosaic-8 RBD-NP constructed from soluble monomeric RBDs (Cohen et al., 2021a) (Fig. 1 B). We used flow cytometry to verify presentation of membrane-bound RBD quartets on the surface of cells transfected with mRNA encoding RBD quartets 4a or 4b by staining with C118, a human mAb that recognizes a conserved RBD epitope present on all eight RBDs (Jette et al., 2021; Robbiani et al., 2020). Flow cytometry showed high and equivalent levels of RBD quartets 4a and 4b on cell surfaces when expressed separately (Fig. S1 A).

We next investigated use of EABR technology, which modifies membrane proteins delivered by mRNA-LNPs to induce self-assembly and budding of eVLPs, resulting in immunogen presentation on cell surfaces and circulating eVLPs (Hoffmann et al., 2023), for delivering quartet RBD sequences. eVLPs are induced by appending a short sequence to the cytoplasmic domain of a membrane protein, which recruits proteins from the host ESCRT pathway that drives budding for many enveloped viruses (Votteler and Sundquist, 2013). Addition of an EABR sequence produces eVLPs for a variety of membrane proteins including SARS-2 spike (Hoffmann et al., 2023), HIV-1 Env (Hoffmann et al., 2023), influenza hemagglutinin (Zhang et al., 2025), RSV fusion protein (Chai et al., 2025), Andes virus glycoprotein (Guo et al., 2025), CCR5 (Hoffmann et al., 2023), epidermal growth factor receptor (Gonzalez-Magaldi et al., 2025), and MHC class I proteins (Olson et al., 2025). Compared to non-EABR RBD quartets, cells expressing EABR-modified RBD quartets showed lower cell surface expression (Fig. S1 A), which presumably results from a subset of cell surface proteins being incorporated into released eVLPs.

To evaluate whether the RBD quartet 4a-EABR and 4b-EABR constructs induced secretion of RBD quartet-displaying eVLPs, we examined eVLPs purified from supernatants of quartet 4a- or 4b–transfected cells. ELISAs were consistent with efficient eVLP formation for EABR mRNAs, showing high quartet 4a or 4b protein levels in purified eVLP samples for EABR, but not non-EABR, quartets (Fig. S1 B). We also used Abs specific for quartet 4a (anti-Rs4081 RBD) (Fan et al., 2025b) and 4b (anti-WIV1 RBD) (Fan et al., 2022) to examine co-transfected cells by flow cytometry, demonstrating that RBD quartets 4a and 4b were co-expressed at individual cell surfaces (Fig. S1 C). To confirm that both quartets were incorporated into the same eVLP, we used a sandwich ELISA to probe eVLPs from an RBD quartet 4a-EABR and 4b-EABR co-transfection. eVLPs were purified, captured with anti-WIV1 RBD Ab (quartet 4b), and detected with anti-Rs4081 RBD Ab (quartet 4a), verifying the presence of both quartets on dual quartet eVLPs (Fig. S1 D). No binding was detected for samples prepared in the same way from cells co-transfected with non-EABR versions of the quartets or cells only transfected with a single quartet (Fig. S1 D). These data show that mRNA-encoded RBD quartets are co-expressed on transfected cells and that the two EABR-modified quartets are co-displayed on eVLPs.

### Dual quartet RBD mRNA immunogens elicit cross-reactive Abs

We next compared Ab responses in mice immunized with mRNA-encoded dual quartets to immunizations with protein-based dual quartet RBD-NPs or with purified dual quartet RBD-eVLPs. mRNAs encoding RBD quartets 4a and 4b were co-formulated into lipid nanoparticles (LNPs) to generate dual quartet immunogens (dual quartet RBD-mRNA and dual quartet RBD-EABR-mRNA), which were each injected intramuscularly at RNA doses of 1.5 µg or 0.5 µg (Fig. 2 A, Fig. S2 A). We also included cohorts of mice that were immunized with 5 µg of dual quartet RBD-NPs (calculated based on RBD content) in the presence of adjuvant (Fig. 2 A) and cohorts immunized with adjuvanted purified RBD-eVLPs (dual quartet RBD-eVLP or SARS-2 RBD-eVLP; 5 µg RBD content per dose), the latter representing a homotypic immunogen control presenting only one type of RBD (Fig. 1 C).

**Figure 2.**
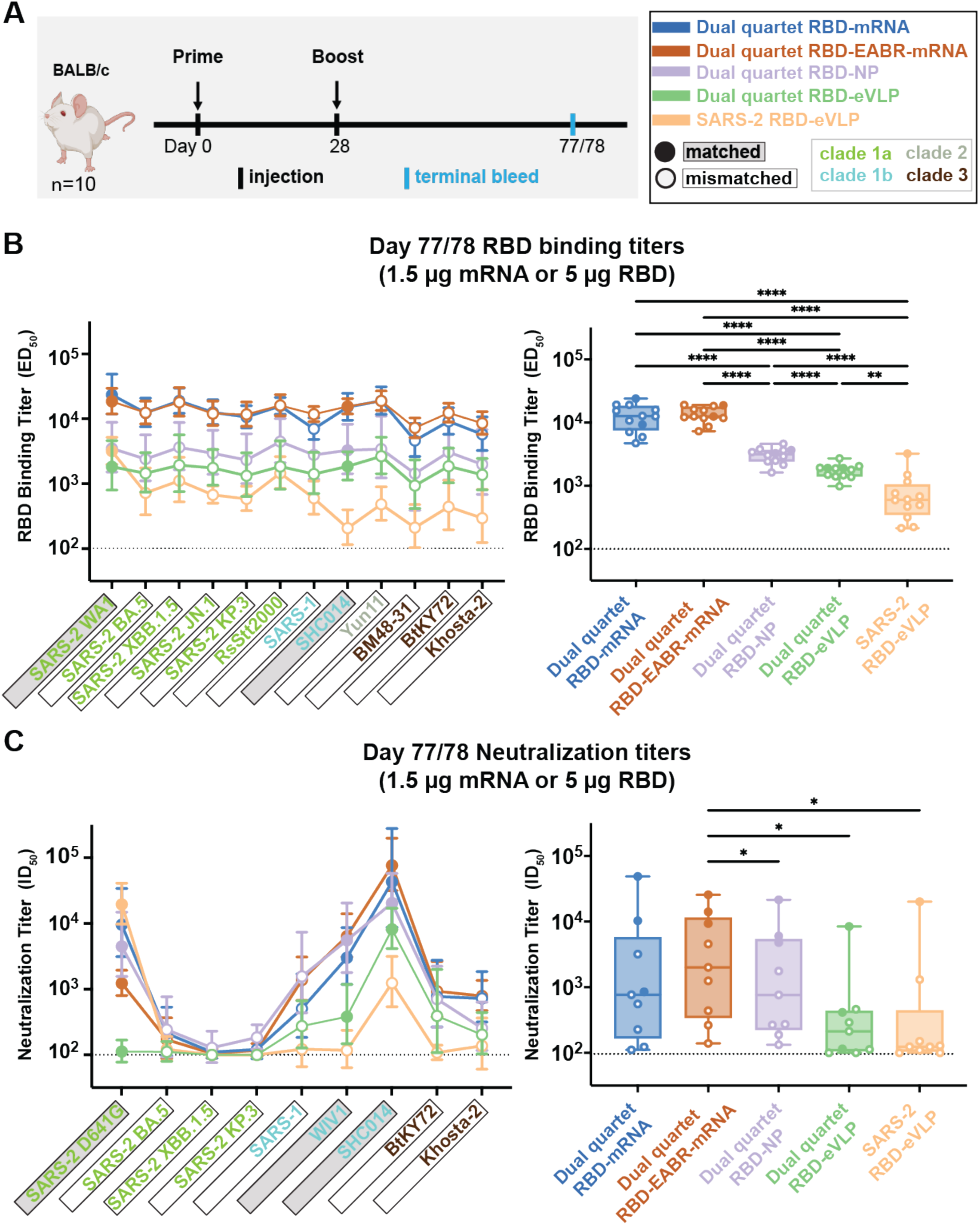
Dual quartet RBD mRNA immunogens elicit cross-reactive Abs. ELISA and neutralization results for terminal bleed serum samples (see also Fig. S2). (A) Immunization regimen. Left: Mice were primed at day 0, boosted at day 28, and samples were collected from a terminal bleed at day 77 or 78. Right: Colors used to identify immunization cohorts and symbols indicating a sarbecovirus antigen or pseudovirus that is matched (filled in data points; gray shading around name) or mismatched (unfilled data points; black outline around name). Sarbecovirus strain names are colored in panels B and C according to clade. (B,C) Results for RBD-binding ELISAs (panel B) and pseudovirus neutralization assays (panel C) for serum samples from day 77 or 78. Dashed horizontal lines indicate detection limits for assays. Left: Geomeans of RBD-binding ELISA half-maximal effective dilution (ED_50_) values (panel B) or neutralization half-maximal inhibitory dilution (ID_50_) values (panel C) across a panel of viral antigens or pseudoviruses for sera from animals in each cohort (geomeans are plotted as symbols with geometric standard deviations indicated by error bars). Mean ED_50_ or ID_50_ values across antigens are connected by colored lines corresponding to the immunization cohort. Right: Box and whisker plots of geomean responses (ELISA ED_50_s in panel B; neutralization ID_50_s in panel C) with individual data points representing geomean responses across n=10 sera to a single antigen or pseudovirus. Geomean titers were compared pairwise between immunization cohorts by Tukey’s multiple comparison test with the Geisser-Greenhouse correction (as calculated by GraphPad Prism). Significant differences between cohorts are indicated by asterisks: p<0.05 = *, p<0.01 = **, p<0.001 = ***, p<0.0001 = ****.

Since binding and neutralizing Ab responses correlate with protection in humans and animals vaccinated with COVID-19 mRNA vaccines (Corbett et al., 2021; Gilbert et al., 2022; Goldblatt et al., 2022), we analyzed immunized mouse sera by ELISA and pseudovirus assays (Schmidt et al., 2020) to evaluate Ab binding responses and neutralization, respectively (Fig. 2, B and C; Fig. S2 B). When comparing mRNA-based to protein-based immunogens, we used a 1.5 µg mRNA dose and a 5 µg (RBD content) dose for the protein-based immunogens (Fig. 2, B and C), representing standard doses in mice (Cohen et al., 2021a; Corbett et al., 2020; Hoffmann et al., 2023). We also compared the mRNA and mRNA-EABR immunogens using a 0.5 µg mRNA dose to investigate potential dose-sparing effects (Fig. S2 B). To evaluate sarbecovirus strain-specific differences in Ab binding and neutralization properties, we displayed data to show responses to individual sarbecovirus antigens (Fig. 2, B and C; left panels) and as overall geometric mean (geomean) responses elicited by different cohorts across evaluated antigens (Fig. 2, B and C; right panels).

Highest mean binding titers for sera across a panel of matched and mismatched sarbecovirus RBDs were found for dual quartet RBD-mRNA and dual quartet RBD-EABR-mRNA cohorts (Fig. 2 B, left). Thus, genetically-encoded RBD quartets delivered by a 1.5 µg dose of mRNA-LNPs were significantly more effective at eliciting cross-reactive RBD-binding Abs across a panel of sarbecovirus RBDs than 5 µg of the dual quartet RBD-NP protein-based immunogen (Fig. 2 B, right; p<0.0001 for comparisons of dual quartet RBD-mRNA and dual quartet RBD-EABR-mRNA). In addition, both dual quartet RBD-mRNA and dual quartet RBD-EABR-mRNA exhibited significantly higher geomean binding titers than the protein-based dual quartet RBD-eVLPs and homotypic SARS-2 RBD-eVLPs (Fig. 2 B, right; p<0.0001 for all comparisons of mRNA-LNP and protein-based immunogens). SARS-2 RBD-eVLPs exhibited significantly lower binding titers across the RBD panel than the dual quartet RBD-NPs and purified dual quartet RBD-eVLPs (Fig. 2 B, right; p<0.0001 for all comparisons except p=0.0057 for dual quartet RBD-eVLP), as expected based on results for homotypic RBD-NPs (Cohen et al., 2021a; Cohen et al., 2022).

We next compared neutralizing titers across a panel of matched and mismatched sarbecovirus strains (Fig. 2 C, left). When comparing by strain between the five immunization cohorts, dual quartet RBD-mRNA, dual quartet RBD-EABR-mRNA, and dual quartet RBD-NP exhibited the highest titers across the pseudovirus panel except for neutralization of SARS-2 D614G, where homotypic SARS-2 RBD-eVLPs elicited slightly higher titers (Fig. 2 C, left). Geomean neutralization titers for the five cohorts (Fig. 2 C, right) were more similar to each other than the mean binding titers (Fig. 2 B, right) but dual quartet RBD-EABR-mRNA induced significantly higher mean neutralizing titers than dual quartet RBD-NP (p=0.0152), dual quartet RBD-eVLP (p=0.0152), and SARS-2 RBD-eVLP (p=0.0470).

We also measured binding and neutralization titers elicited by 0.5 µg doses of dual quartet RBD-mRNA and dual quartet RBD-EABR-mRNA (Fig. S2, A and B). Binding titers against the RBDs tested were not statistically different for the two mRNA immunogens, except for binding to the BM48-31 RBD, in which case dual quartet RBD-EABR-mRNA induced significantly higher binding than dual quartet RBD-mRNA (Fig. S2 B, left; p=0.0089). Neutralization titers were significantly different for three viruses: SARS-2 D614G (titers higher for dual quartet RBD-mRNA; p=0.00013) and for SARS-1 and WIV1 (higher for dual quartet RBD-EABR-mRNA; p=0.00024 and p<0.0001, respectively) (Fig. S2 B, right panel). These results are consistent with more potent neutralization of non-SARS-2 viruses induced by dual quartet RBD-EABR-mRNA compared with dual quartet RBD-mRNA at the 1.5 µg dose (Fig. 2 C).

### mRNA-encoded dual quartets elicit potent T cell responses

T cell responses to the various immunogens were evaluated on day 77/78 in immunized mouse splenocytes by ELISpot (Ranieri et al., 2014). Cytokines (IFN-γ produced by CD4+ T helper 1 (T_H_1) cells and cytotoxic CD8+ T cells; IL-4 produced by CD4+ T helper 2 (T_H_2) and T follicular helper cells (Leehan and Koelsch, 2015)) were measured after stimulation with a pool of SARS-2 spike peptides.

Dual quartet RBD-mRNA and dual quartet RBD-EABR-mRNA cohorts exhibited potent IFN-γ responses, consistent with activation of RBD-specific cytotoxic CD8+ T cells, whereas low/undetectable IFN-γ responses were found for the protein-based dual quartet RBD-NP, dual quartet RBD-eVLP, and SARS-2 RBD-eVLP cohort, consistent with previous findings for mice immunized with purified spike-EABR eVLPs (Hoffmann et al., 2023) and the fact that mRNA-LNPs, but not protein immunogens, promote intracellular expression of antigens that serve as sources for peptides loaded onto MHC class I proteins (Rock et al., 2016). We found strong IL-4 responses for both the mRNA and protein-based immunogen cohorts (Fig. 3).

**Figure 3.**
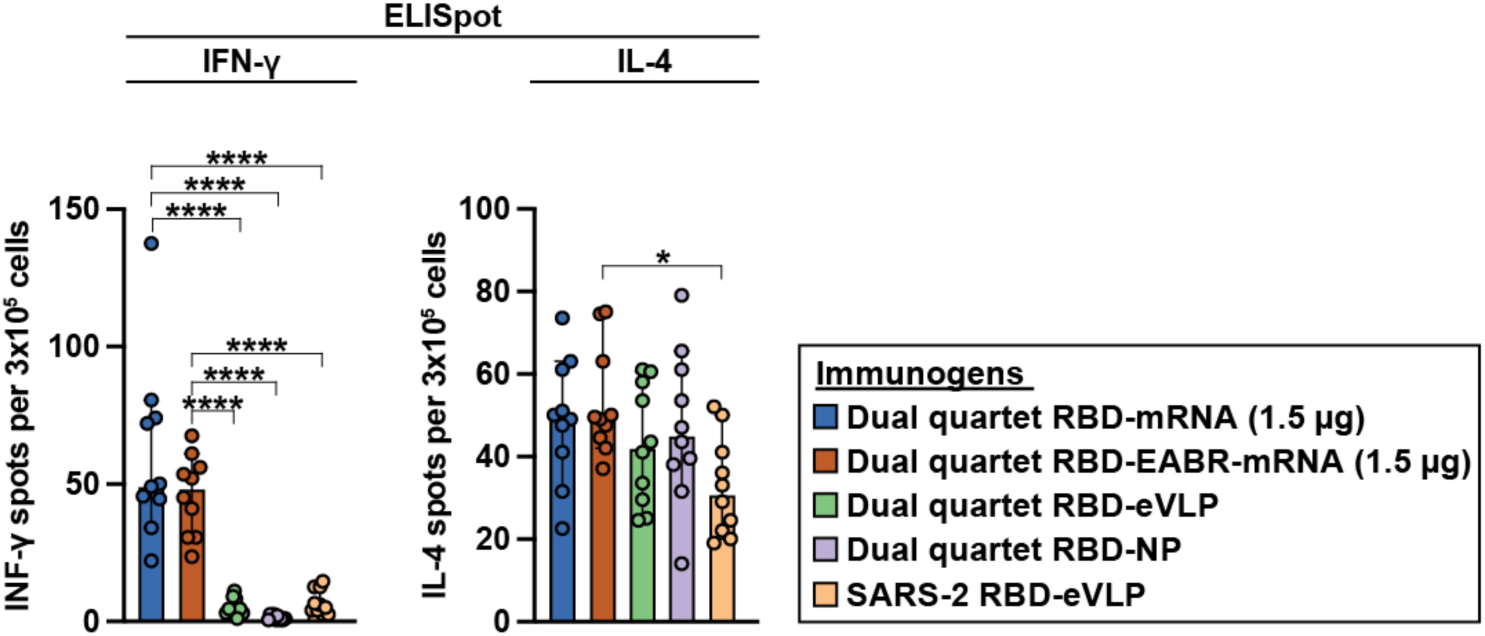
mRNA-encoded dual quartets elicit potent T cell responses. ELISpot assay data for SARS-2 RBD-specific IFN-γ (left) and IL-4 (right) responses of splenocytes from BALB/c mice that were immunized with the indicated immunogens (immunization regimen in Fig. 2 A). Results are shown as spots per 3×10^5^ cells for individual mice (colored circles) presented as the median (bars) and standard deviation (horizontal lines). Cohorts were compared by Tukey’s multiple comparison test calculated by GraphPad Prism. Significant differences between cohorts linked by horizontal lines are indicated by asterisks: p<0.05 = *, p<0.0001 = ****.

Thus, mRNA-based mosaic-8 dual RBD quartet immunizations induce superior (INF-γ) or equivalent (IL-4) release of cytokines compared with protein-based mosaic-8 immunogens.

### mRNA-encoded and protein-based dual quartets elicit Abs targeting similar epitopes

To map RBD epitopes targeted by Abs elicited by different vaccine modalities, we used DMS to evaluate effects of all possible RBD amino acid changes on binding to polyclonal serum Abs (Greaney et al., 2021a; Greaney et al., 2021b; Greaney et al., 2021c; Greaney et al., 2021d; Starr et al., 2021a; Starr et al., 2021b; Starr et al., 2020). We classified RBD residues by epitopes defined by structural properties and conservation/variability across sarbecoviruses: class 1 and class 2 RBD epitopes exhibit more, and class 4, class 1/4, and portions of class 3 and class 5 epitopes exhibit less, sequence variability across sarbecoviruses and SARS-2 variants (Barnes et al., 2020; Cui et al., 2024; Jensen et al., 2023; Jette et al., 2021) (Fig. 4 A). We analyzed serum using yeast display libraries encoding RBDs derived from SARS-2 WA1 (matched) and SARS-1 (mismatched) spikes.

**Figure 4.**
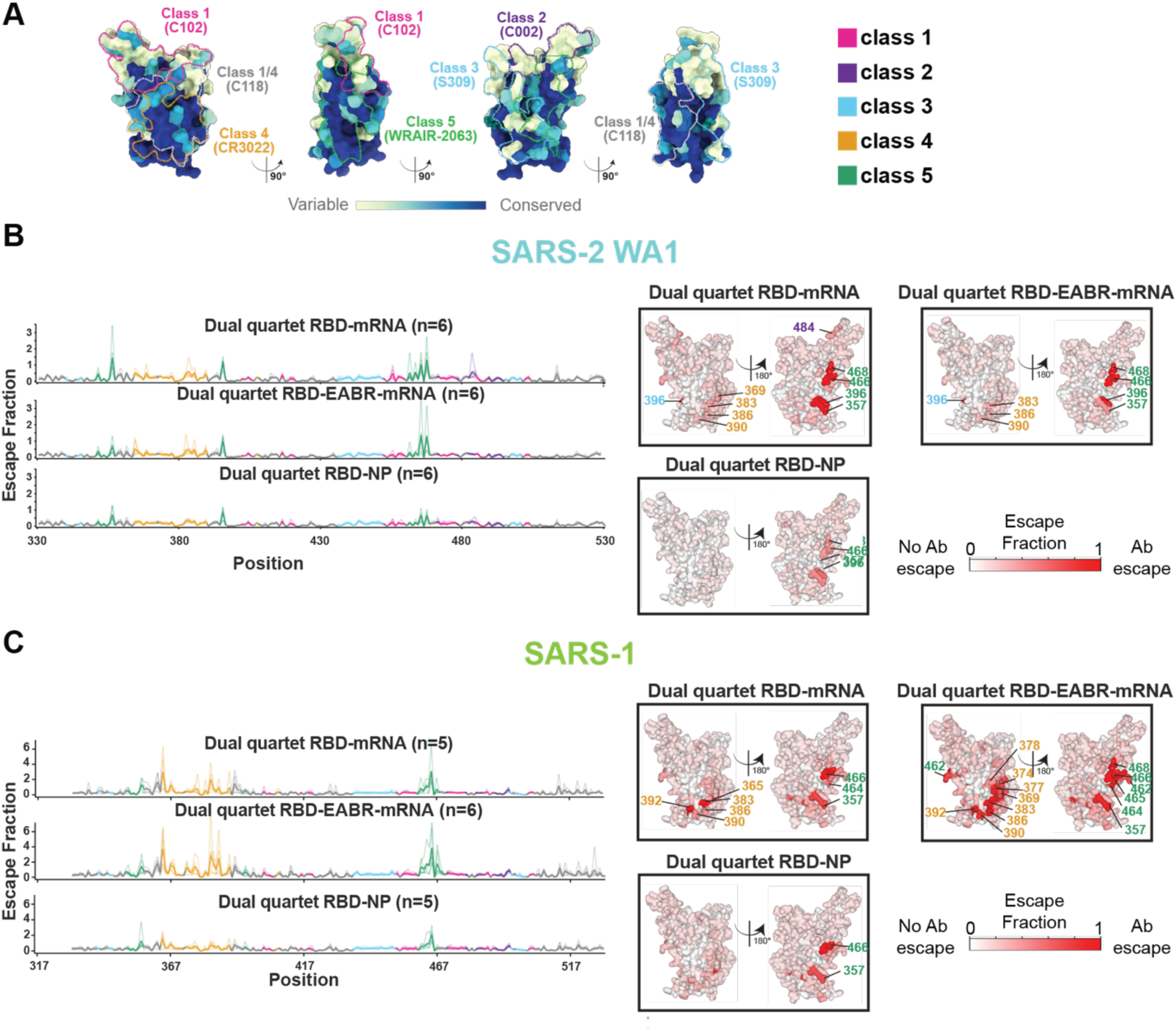
Dual quartet RBD-mRNA immunogens elicit Ab responses against conserved class 4 and class 5 RBD epitopes. (A) Sequence conservation of 16 sarbecovirus RBDs calculated using ConSurf (Landau et al., 2005) shown on a surface representation of SARS-2 RBD (PDB 7BZ5). Class 1, 2, 3, 4, 1/4, and 5 anti-RBD Ab epitopes (Barnes et al., 2020; Cui et al., 2024; Jensen et al., 2023; Jette et al., 2021) are outlined in dots in different colors using information from representative structures of Abs bound to SARS-2 spike or RBD (C102: PDB 7K8M; C002: PDB 7K8T, S309: PDB 7JX3; CR3022: PDB 7LOP; C118: PDB 7RKV; WRAIR-2063: PDB 8EOO). (b,c) Left: Line plots for DMS results using a SARS-2 WA1 RBD library (B) and SARS-1 library (C) from the indicated number of mice immunized with the immunogens listed above each line plot. X-axis: RBD residue number. Y-axis: sum of the Ab escape of all mutations at a site (larger numbers = more Ab escape). Each line represents one antiserum with thick lines showing the average across the n = 5 or 6 sera in each group. Lines are colored according to RBD epitopes in panel A. Right: Average site-total Ab escape calculated for results from n = 5 or 6 serum samples for the indicated RBD yeast display libraries. Mice were immunized with the immunogens listed, and results were mapped to the highlighted residues on the surface of the SARS-2 WA1 RBD (PDB 6M0J). Gray indicates no escape; a gradient of red represents increasing degree of escape. Residue numbers show sites with the most escape with font colors representing different RBD epitopes (defined in panel A; class 1/4 residues are colored with fonts corresponding to class 1 or class 4 residues). The same data are shown in Table 1 and in Figs. S3 and S4 (line and logo plots for SARS-2 WA1 and SARS-1 RBD libraries, respectively).

**Table 1.**
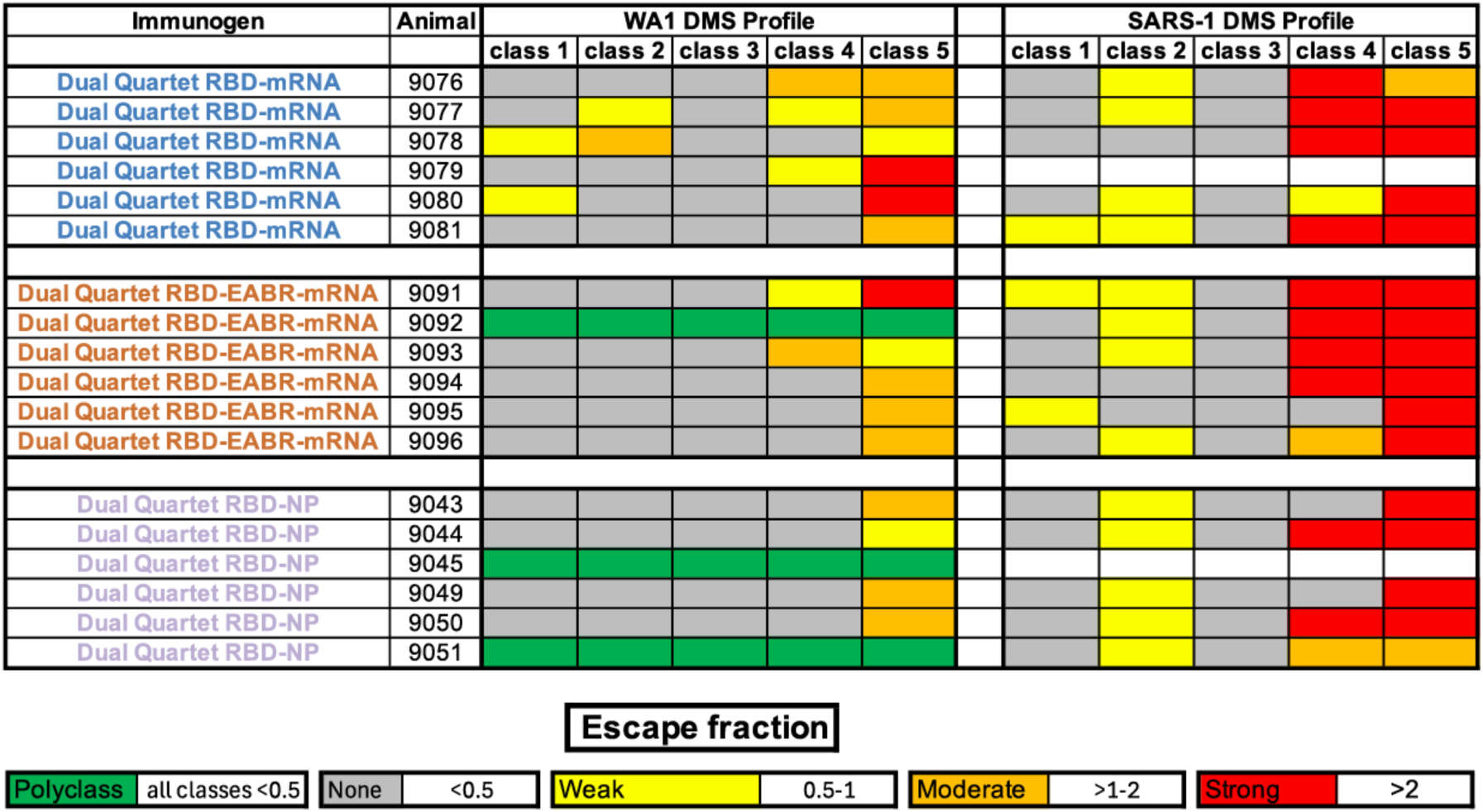
Summary of DMS data showing elicitation of strong Ab responses against class 4 and class 5 RBD epitopes by dual quartet RBD-mRNA immunogens. Results for individual mice are shown with ID numbers indicated. DMS was conducted using a SARS-2 WA1 RBD library (left) or a SARS-1 RBD library (right). DMS profiles were classified as polyclass, weak, moderate, or strong as indicated by the maximum escape fraction for any RBD position in each RBD epitope class. Colors and different designations for epitopes were assigned as indicated at the bottom (white entries for animals 9079 and 9045 indicate that no data were available). See also Fig. 4 and Figs. S3 and S4.

DMS showed that the epitopes targeted in both RBD libraries by the mRNA-encoded dual quartet RBD immunogens were similar overall to each other and to epitopes targeted by the protein-based dual quartet RBD-NP immunogen (Fig. 4, B and C; Table 1; Figs. S3 and S4). The dual quartet immunogens, like mosaic-8 RBD-NPs (Cohen et al., 2024; Cohen et al., 2022), did not primarily elicit Abs against the immunodominant and variable class 1 and class 2 RBD epitopes, as observed for Abs elicited by homotypic SARS-2 RBD NPs (Cohen et al., 2024; Cohen et al., 2022) and spike-based mRNA vaccines (Bangaru et al., 2025). Instead, elicited Abs primarily targeted class 4 and class 5 RBD epitopes, except for the Abs raised by immunization with dual quartet RBD-NPs, where class 5 was the predominant targeted epitope.

Although dual quartet RBD immunizations resulted in a mixed class 4/class 5 RBD response when evaluated against the SARS-2 WA1 and SARS-1 RBD libraries (Fig. 4, B and C; Table 1; Figs. S3 and S4), there were notable differences. For example, both RNA-based immunogens elicited similar class 4 and class 5 RBD responses, but dual quartet RBD-mRNA sera showed more of a class 1/class 2 RBD response than dual quartet RBD-EABR-mRNA sera against the SARS-2 WA1 RBD library (Fig. 4 B and Table 1), perhaps explaining why dual quartet RBD-mRNA elicited higher neutralization titers against the matched SARS-2 D614G pseudovirus (Fig. 2 C and Fig. S2 B). Against the SARS-1 RBD library, each of the dual quartet immunogens induced strong class 4 and class 5 responses. In addition, weak escape by Abs against the variable class 1 and class 2 RBD epitopes was seen for dual quartet RBD-mRNA, dual quartet RBD-EABR-mRNA, and dual quartet RBD-NP (Table 1). Finally, one mouse in the dual quartet RBD-EABR-mRNA group and two in the dual quartet RBD-NP group elicited “polyclass” responses (defined as weak escape profiles that lacked clear features, which we interpret as resulting from sera containing multiple classes of anti-RBD Abs (Cohen et al., 2024), consistent with the ability of these immunogens to induce Abs against multiple RBD epitopes.

We conclude that converting dual quartet RBD immunogens into an mRNA format retains the ability of mosaic-8 vaccines to elicit Abs mainly against more conserved RBD regions, as previously observed in DMS experiments comparing protein NPs presenting dual RBD quartets or eight individual RBDs (Hills et al., 2024).

### mRNA-encoded dual quartet RBD immunogens elicit balanced IgG2a/IgG1 responses

IgG subclasses induce distinct Fc effector functions, including opsonization, phagocytosis, and Ab-dependent cell-mediated cytotoxicity, through differential binding of Fcs to Fc gamma receptors (FcγRs) (Crescioli et al., 2025); e.g., mouse IgG1, IgG2a, and IgG2b bind the activating receptor FcγR3 and the inhibitory receptor FcγR2b, IgG2a and IgG2b bind the activating receptor FcγR4, and mouse IgG3 does not bind detectably to these FcγRs (Bruhns and Jonsson, 2015). FcγR3 is more widely expressed than FcγR4, with FcγR4 mainly expressed on monocytes/macrophages and neutrophils and FcγR3 found on those cells plus dendritic cells, natural killer cells, basophils, mast cells, and eosinophils (Bruhns and Jonsson, 2015). In immunized mice, IgG2a is associated with T_H_1-type immune responses, effective activation of complement, and protective interactions with FcγRs, which are critical functions for clearing viruses (Hwang et al., 2021). IgG1, although usually efficient in neutralizing viruses, is associated with T_H_2-type responses thought to be less effective than T_H_1 responses in combatting viruses (Aleebrahim-Dehkordi et al., 2022).

We used systems serology (Ackerman et al., 2017) to evaluate IgG distributions elicited by mRNA versus protein immunogens by comparing binding of IgG1, IgG2a, IgG2b, IgG3, FcγR2b-, FcγR3-, or FcγR4-binding IgGs, and total IgG (IgGs isolated using a pan-IgG Ab) elicited by the five immunogens to a panel of spike trimers and RBDs derived from SARS-2 variants and other sarbecoviruses. Results show elicited responses from individual mice to each antigen (Fig. 5) and geomean responses across antigens for each immunogen (Fig. S5).

**Figure 5.**
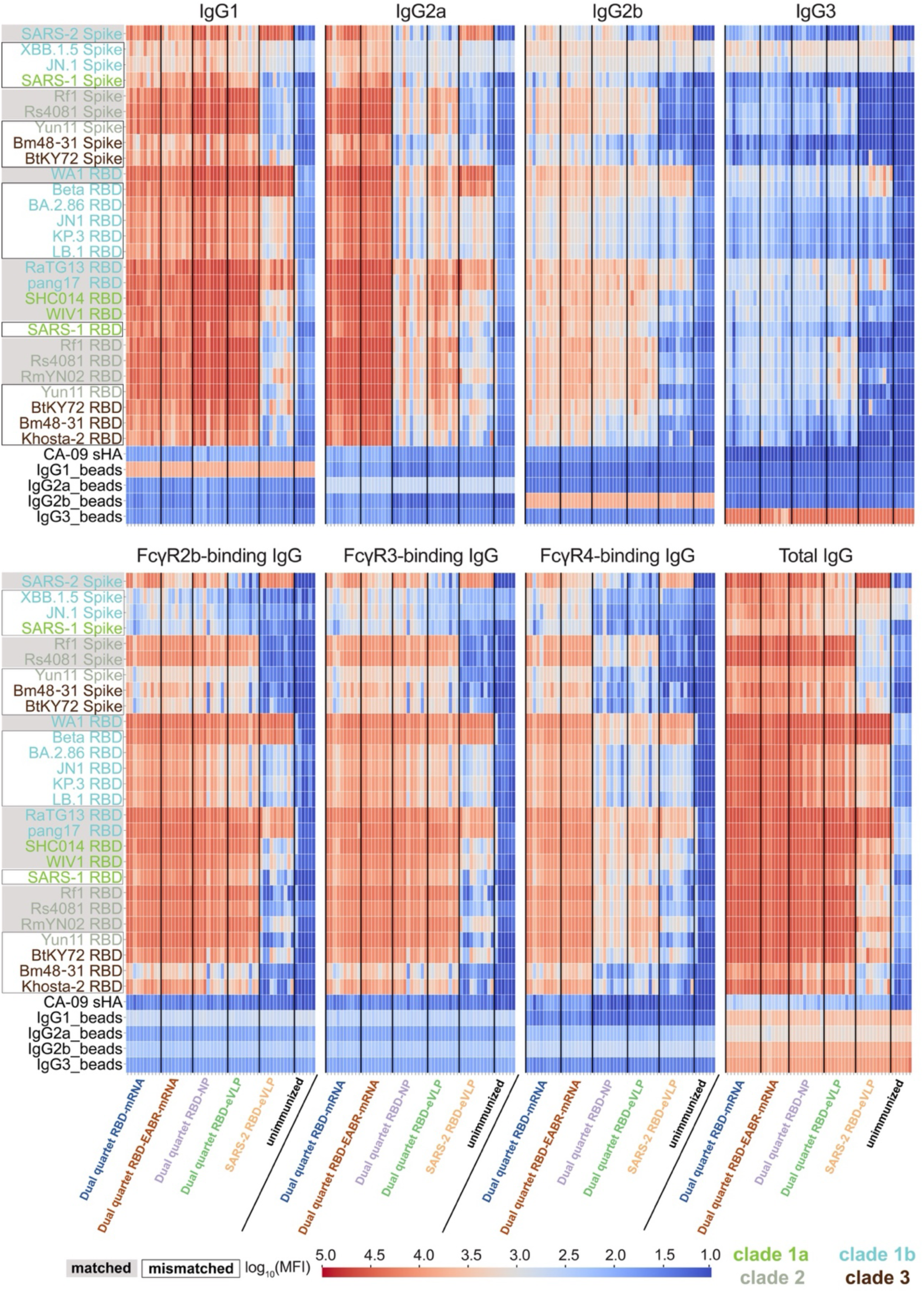
mRNA-encoded dual quartet RBD immunogens elicit balanced IgG subclass and potent FcγR-binding responses. MFI = median fluorescent intensity. Binding of IgG1, IgG2a, IgG2b, IgG3, FcγR2b-binding IgGs, FcγR3-binding IgGs, FcγR4-binding IgGs, and total IgG to the indicated spikes, RBDs, or non-sarbecovirus control proteins. Antigen names (y-axes) are colored according to spike or RBD clades. Matched antigens are indicated with gray shading around the name, and mismatched antigens are indicated with a black outline around the name. Each column represents binding data from an individual mouse, and immunization cohorts are separated by a vertical black line. A soluble influenza hemagglutinin trimer (CA-09 sHA) was used as a control to evaluate non-specific binding. See also Fig. S5.

Both dual quartet RBD-mRNA immunogens elicited high levels of IgG1 and IgG2a against sarbecovirus spikes and RBDs, with somewhat higher levels of IgG2a than IgG1 (Fig. 5 and Fig. S5), demonstrating a desirable mixture of IgGs for an anti-viral vaccine of induced T_H_1 and T_H_2 responses. In addition, the mRNA-based immunogens elicited significantly higher IgG2a and FcγR4-binding IgG responses than the protein-based immunogens (p<0.0001) (Fig. S5). Dual quartet RBD-NPs and dual quartet RBD-eVLPs elicited higher levels of IgG1 than IgG2a, consistent with their administration with Addavax adjuvant, which enhances T_H_2-biased immune responses (Han et al., 2019). Dual quartet RBD-eVLPs also elicited significantly higher IgG2a than dual quartet RBD-NPs (p<0.0001). Interestingly, dual quartet RBD-EABR-mRNA elicited significantly higher IgG2b (p<0.0001), as well as FcyR3-(p=0.0013) and FcyR4-(p=0.0021) binding IgGs, than dual quartet RBD-mRNA. Consistent with demonstrations of more limited sarbecovirus recognition properties of Abs raised in response to homotypic RBD-NPs (Cohen et al., 2021a; Cohen et al., 2024; Cohen et al., 2022; Fan et al., 2025b), SARS-2 RBD-eVLP elicited IgGs mainly against SARS-2 and closely related RBDs (Fig. 5).

Thus, mRNA versions of dual RBD quartets elicit balanced IgG responses that broadly recognize sarbecovirus antigens, arguing for their likely efficacy to protect against diverse sarbecoviruses.

### Systems Serology-Polyclonal Epitope Mapping (SySPEM)

To further explore potential differences in RBD recognition properties by different IgG subclasses within polyclonal Ab samples, we combined systems serology with epitope mapping in a new methodology we call Systems Serology-Polyclonal Epitope Mapping (SySPEM) (Fig. S6), by analogy to Electron Microscopy-based Polyclonal Epitope Mapping (EMPEM) (Turner et al., 2023). For epitope mapping by SySPEM, we created epitope knock-out (KO) mutants in the SARS-2 WA1 RBD by introducing one or more potential N-linked glycosylation sites (PNGSs) to direct addition of N-glycan(s) in RBD epitopes that are recognized by either class 1 and class 2, class 2 only, class 3, class 1/4, class 4, or class 5 anti-RBD mAbs (Fig. S7 A). Addition of N-glycan(s) at introduced PNGS(s) was verified by SDS-PAGE of purified RBD KO proteins, and correct folding and epitope blocking were verified using a panel of control mAbs (Fig. S7, B and C). We measured binding of total polyclonal IgG and IgG subsets to unmodified (wildtype; wt) WA1 RBD and to each RBD KO mutant, defining a SySPEM score for an IgG sample/epitope pair as [1.0 – (RBD KO binding / RBD wt binding) x 100]; thus, higher SySPEM scores indicate increased binding by a particular IgG class to that epitope (Fig. S6). We present SySPEM results in three formats to visualize epitope recognition across immunogens and Ab subclasses: (*i*) Median SySPEM scores from individual mice divided into distinct RBD epitopes recognized by different IgG classes with statistical comparisons between immunogen cohorts (Fig. 6). This comparison highlights differences in epitope recognition across immunogen cohorts. (*ii*) Median SySPEM scores for each IgG class within a given immunogen cohort with statistical comparisons between targeted epitopes (Fig. S8). This shows the epitope distribution for a given immunogen/IgG class pair, allowing for epitope profile comparisons between different immunogens, analogous to a DMS profile (e.g., Fig. 4). (*iii*) Uniform Manifold Approximation and Projection (UMAP) (McInnes et al., 2020) format to reduce the data set dimensions (Fig. 7). These figures provide hierarchical views of the SySPEM data: statistical comparisons across immunogens (Fig. 6), detailed within-cohort analyses (Fig. S8), and global similarity mapping of Ab recognition patterns (Fig. 7).

**Figure 6.**
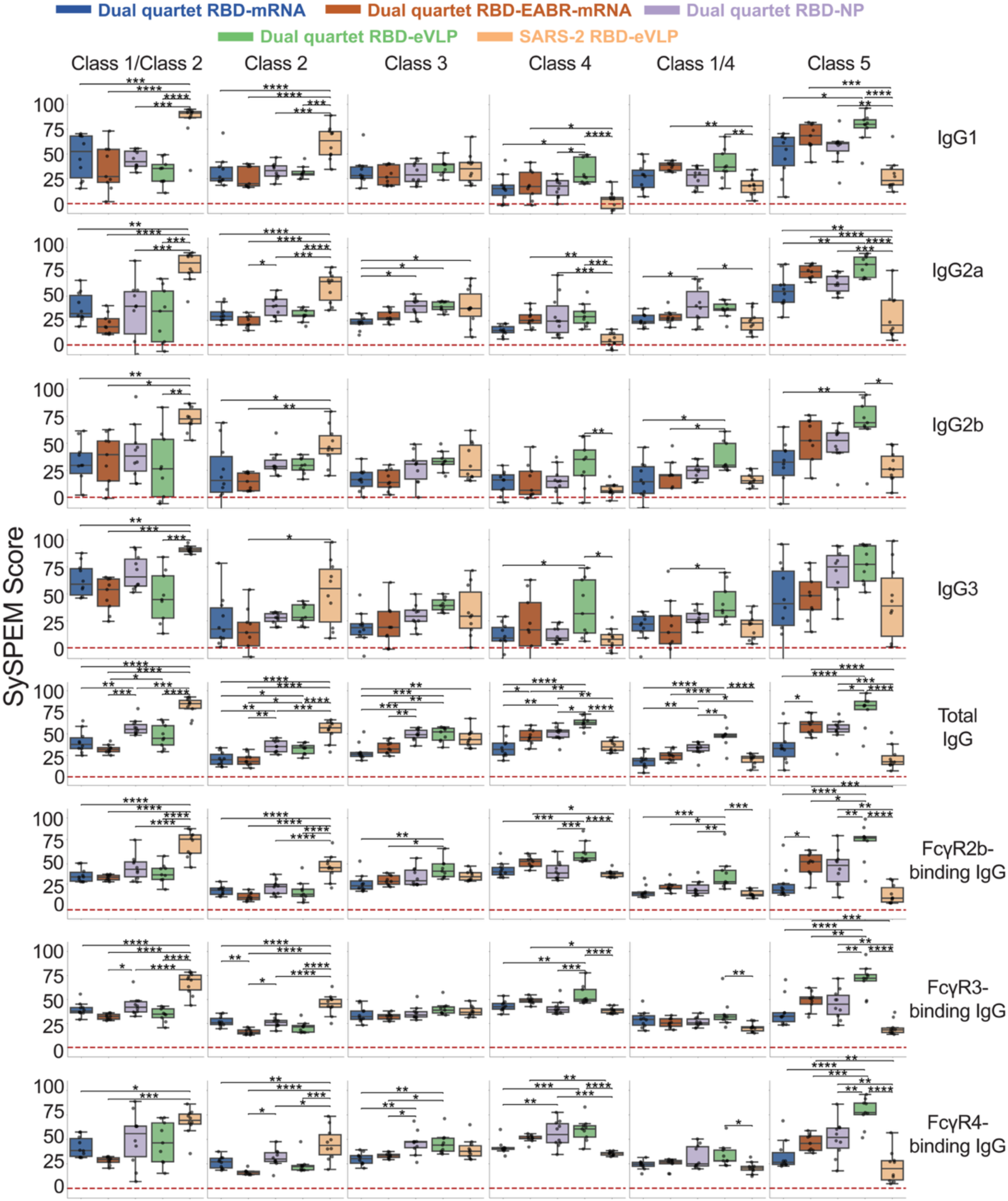
SySPEM score comparisons show differences in epitope recognition across immunogen cohorts. SySPEM scores from individual mice were determined for RBD epitopes (columns) recognized by different IgG classes (rows, IgG subclass plus different FcγR-binding IgGs) with statistical comparisons between immunogen cohorts (colors). A SySPEM value of 0 indicates that none of the IgGs in that sample were affected by the glycan addition and therefore the sample did not contain IgGs that recognize that epitope, and a SySPEM value of 100 indicates that all IgGs in that sample recognized that epitope (Fig. S6). A SySPEM value between 0 and 100 indicates the proportion of IgGs in a sample that recognized the epitope that was blocked by glycan addition in the RBD KO mutant. Box and whisker plots of SySPEM scores with individual data points representing one mouse are shown. Significant differences between cohorts were calculated using Tukey’s HSD posthoc test and linked by vertical lines indicated by asterisks: p<0.05 = *, p<0.01 = **, p<0.001 = ***, p<0.0001 = ****. See also Fig. 7 and Fig. S8.

**Figure 7.**
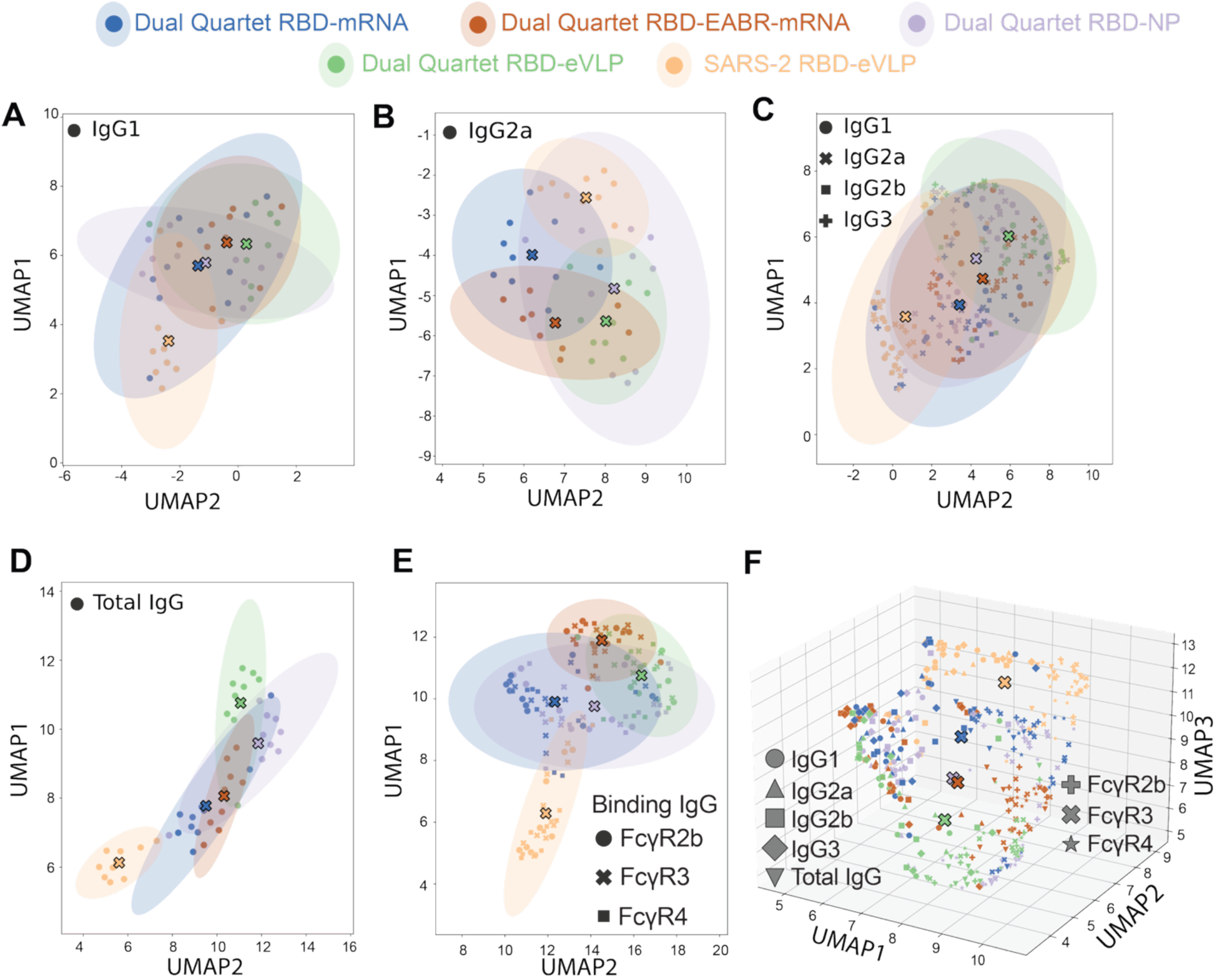
SySPEM UMAP analyses. Uniform Manifold Approximation and Projection (UMAP) (McInnes et al., 2020) was used to project multi-dimensional SySPEM scores into two (panels A-E) or three (panel F) dimensions for (A) IgG1 (B) IgG2a, (C) IgG1, IgG2a, IgG2b, and IgG3, (D) Total IgG, (E) FcγR-binding IgGs, and (F) Total IgG, FcγR-binding IgGs, and IgG subclasses. Samples with similar Ab–epitope recognition profiles cluster together, with colored ellipses indicating the variance within each immunogen group and centroid markers (large Xs) showing the group mean. (F) Three-dimensional UMAP projection for all IgG subclasses and FcγR-binding IgGs with centroid markers (large Xs). See also Fig. 6, Figs. S6 and S8.

SySPEM scores validated the mapping approach using RBD KO mutants by showing significantly higher scores for total IgGs elicited by homotypic SARS-2 RBD-eVLPs than by other immunogens against variable epitopes (class 1/class 2, class 2) and little to no targeting of more conserved epitopes (class 4, class 1/4, class 5) (Fig. 6), as previously observed by DMS for homotypic SARS-2 RBD-NPs (Cohen et al., 2022).

For IgG1, the mRNA- and protein-based dual quartet immunogens showed the highest class 5 SySPEM scores, with only dual quartet RBD-mRNA exhibiting a score that was not significantly higher than that of homotypic SARS-2 RBD-eVLPs (Fig. 6). In addition, dual quartet RBD-eVLPs and dual quartet RBD-EABR-mRNA exhibited class 4 and class 1/4 IgG1 SySPEM scores that were significantly higher than respective scores for SARS-2 RBD-eVLPs. Similarly, for IgG2a, the dual quartet immunogens all elicited significantly higher class 5 responses than homotypic SARS-2 RBD-eVLP, with both dual quartet RBD-EABR-mRNA and dual quartet RBD-eVLP featuring the highest class 5 SySPEM scores. Dual quartet RBD-EABR-mRNA, dual quartet RBD-NP, and dual quartet RBD-eVLP also elicited higher class 4 SySPEM scores than SARS-2 RBD-eVLP. Although not significant, dual quartet RBD-mRNA elicited a higher class 1/class 2 SySPEM score than dual quartet RBD-EABR-mRNA, and the opposite trend was observed for class 5. These differences might explain the more potent SARS-2 D614G neutralization elicited by dual quartet RBD-mRNA compared to dual quartet RBD-EABR-mRNA (Fig. 2 C), since class 1 and class 2 anti-RBD Abs are potent against SARS-2 WA1 (Barnes et al., 2020). However, the combination of higher class 5 and lower class 1/class 2 responses elicited by the dual quartet RBD-EABR-mRNA immunogen represents a more favorable epitope targeting profile for broad pan-sarbecovirus vaccine efforts. For IgG2b and IgG3, SySPEM scores were more similar across the different RBD epitope categories, although dual quartet RBD-eVLP showed significantly higher class 4 and class 5 SySPEM scores than SARS-2 RBD-eVLP.

For total IgG, dual quartet RBD-EABR-mRNA and dual quartet RBD-NP showed higher class 5 SySPEM scores than SARS-2 RBD-eVLP, with dual quartet RBD-eVLP showing the highest scores, and dual quartet RBD-mRNA showing class 5 scores that were not significantly higher than those exhibited by SARS-2 RBD-eVLP (Fig. 6). Dual quartet RBD-EABR-mRNA exhibited significantly higher class 4 and class 5 SySPEM scores for total IgG than dual quartet RBD-mRNA, likely driven by the stronger class 4 and class 5 targeting by IgG2a for this comparison (Fig. 6). Class 4 responses were higher for dual quartet RBD-EABR-mRNA, dual quartet RBD-NP, and dual quartet RBD-eVLP all exhibiting significantly higher class 4 SySPEM scores than SARS-2 RBD-eVLP, with dual quartet RBD-eVLP showing the highest class 4, 1/4, and 5 scores.

FcγR2b- and FcγR3-binding IgG SySPEM scores were similar across the RBD epitopes, with dual quartet RBD-EABR-mRNA, dual quartet RBD-NP, and dual quartet RBD-eVLP all exhibiting significantly higher class 5 SySPEM scores than SARS-2 RBD-eVLP (Fig. 6). Dual quartet RBD-eVLP and, to a lesser extent, dual quartet RBD-EABR-mRNA exhibited higher class 4 SySPEM scores than SARS-2 RBD-eVLPs. For FcγR4-binding IgGs, dual quartet RBD-EABR-mRNA, dual quartet RBD-NP, and dual quartet RBD-eVLP showed significantly higher class 4 and class 5 SySPEM scores than SARS-2 RBD-eVLP.

To identify all epitopes recognized by sera elicited by a given immunogen, we displayed the SySPEM results in a format in which SySPEM scores for recognition of individual RBD epitopes by different IgG classes are shown (Fig. S8). These results highlight increased targeting of class 5 RBD epitopes by IgG1, IgG2a, IgG2b, and IgG3 subclasses elicited by the four dual quartet immunogens, but not by homotypic SARS-2 RBD-eVLPs. Displaying the data in this format showed that in some cases for a given immunogen, different IgG subclasses can exhibit distinct epitope profiles, e.g., IgG1 and IgG2a from dual quartet EABR-mRNA sera, where IgG1 showed higher class 1 and class 2 targeting than IgG2a, for which only class 5 targeting was high.

To visualize global patterns of epitope recognition between the different immunogens, we made UMAPs (McInnes et al., 2020) to project epitope profiles into two- or three-dimensional plots (Fig. 7). For 2D UMAPs (Fig. 7, A-E), clusters of points indicate samples that behaved similarly across all epitope knockouts, with ellipses indicating the spread of responses from each immunogen, and centroid markers (indicated by an X) indicating the average epitope mapping profile for each immunogen. When comparing IgG1 (Fig. 7 A), IgG2a (Fig. 7 B), IgG1, IgG2a, IgG2b, and IgG3 (Fig. 7 C), and total IgG (Fig. 7 D), epitope profiles for different immunogens showed similar trends with SARS-2 RBD-eVLP clustering farthest from the four dual quartet immunogens, and the dual quartet immunogens showing more overlap with each other. Dual quartet RBD-mRNA mapped closer to SARS-2 RBD-eVLP in all cases (Fig. 7, A-E), as is especially apparent for IgG2a (Fig. 7 B), total IgG (Fig. 7 D), and the FcγRs (Fig. 7 E). By contrast, dual quartet RBD-EABR-mRNA clustered further away from SARS-2 RBD-eVLP and closer to dual quartet RBD-eVLP. Dual quartet RBD-NP mapped similarly to dual quartet RBD-EABR-mRNA. To further assess differences in epitope recognition between different immunogens, a three-dimensional UMAP plot was generated using all data (Fig. 7 F). Trends in the 3D map were consistent with the 2D UMAPs in that responses to the dual quartet immunogens were distinct from responses to SARS-2 RBD-eVLP and responses to dual quartet RBD-EABR-mRNA tended to map between dual quartet RBD-mRNA and the protein-based dual quartets, consistent with the EABR approach that combines traditional mRNA- and protein-based vaccinations (Hoffmann et al., 2023). In addition, finding that SySPEM scores for dual quartet RBD-mRNA overlapped more with scores for SARS-2 RBD-eVLP than scores for the other dual quartet immunogens is consistent with more class 1/class 2 RBD epitope targeting by dual quartet RBD-mRNA and SARS-2 RBD-eVLP (Table 1).

Overall, the SySPEM studies confirmed targeting of more conserved RBD epitopes by the mRNA-based dual quartet immunogens, particularly the EABR-modified version, indicating successful conversion of the desirable properties of protein-based mosaic-8 immunogens. With detailed analyses, however, we found distinct epitope profiles for IgG subclass and FcγR-binding IgGs: i.e., the same antigens presented via different modalities (mRNA plus and minus EABR sequences and protein antigens on NPs or eVLPs) showed divergent SySPEM epitope profiles. In addition, SySPEM profiling of binding data rationalized differences in neutralization profiles when comparing two similar immunogens presented differently (e.g., dual quartet RBD-EABR-mRNA versus dual quartet RBD-mRNA).

## Discussion

mRNA-based COVID-19 vaccines have been critical for preventing severe disease and death after SARS-2 infection (Wong and Mabbott, 2024), thereby helping keep the worldwide pandemic in check. A recent modification of Moderna’s mRNA-1273 vaccine involved changing the encoded antigen from a SARS-2 spike trimer to a membrane-bound spike N-terminal domain and RBD (Stewart-Jones et al., 2023). The new vaccine, mRNA-1283, exhibited equivalent or superior induced immune responses compared to mRNA-1273 in mice (Stewart-Jones et al., 2023) and humans (Chalkias et al., 2025), thus supporting further development of RBD-based COVID-19 and broader CoV vaccines.

We previously described protein-based RBD-NP immunogens that show promise as a pan-sarbecovirus vaccine (Cohen et al., 2021a; Cohen et al., 2024; Cohen et al., 2022; Fan et al., 2022; Fan et al., 2025b; Hills et al., 2024; Wang et al., 2025). However, manufacturing a protein-based mosaic-8 NP vaccine presents challenges: mosaic-8 RBD-NP requires expression and purification of nine protein components (eight RBDs and a NP) to generate a conjugated RBD-NP, which then requires additional purification and characterization. mRNA vaccines offer the possibility of simpler manufacturing (Pardi et al., 2018) but developing eight individual RBD constructs would also be challenging, especially since mRNA delivery might not achieve co-display of all eight RBDs on cell surfaces and/or eVLPs.

Here, we report an efficient approach to develop an mRNA-encoded version of a broadly cross-reactive mosaic-8 RBD-NP vaccine. The new approach, inspired by protein-based dual quartet RBD-NPs (Hills et al., 2024), involves genetically encoding sarbecovirus RBDs arranged as two membrane-bound quartets, each comprising four tandemly arranged RBD ectodomains. RBD quartets offer the opportunity for optimal Ab induction by displaying a greater number of RBDs per transmembrane region compared with presentation of individual RBDs. The mRNA immunogens described here display dual RBD quartets at cell surfaces (dual quartet RBD-mRNA) or at both the surfaces of cells and secreted eVLPs (dual quartet RBD-EABR-mRNA) when an EABR cytoplasmic domain sequence is included (Hoffmann et al., 2023). Our analyses showed that dual quartet mRNA immunogens elicited equivalent or superior Ab responses, improved FcγR binding of elicited Abs, and increased T cell responses compared with protein-based dual quartet RBD-NPs. In addition, the dual quartet mRNA immunogens induced balanced T_H_1/T_H_2 responses, thereby stimulating both cellular and humoral immune responses. Finally, of critical importance for a potential pan-sarbecovirus vaccine, DMS showed that mRNA-based dual RBD quartet immunogens induced Abs against conserved RBD epitopes, as required for a pan-sarbecovirus vaccine and as observed for protein-based mosaic-8 RBD-NP immunogens (Cohen et al., 2021a; Cohen et al., 2024; Cohen et al., 2022; Fan et al., 2022; Fan et al., 2025b; Hills et al., 2024; Wang et al., 2025).

Despite eliciting high Ab binding titers against both SARS-2 variants and non-SARS-2 sarbecoviruses, neutralization titers for mosaic-8 immunogens (dual quartet RBD mRNAs, dual quartet RBD-NPs, and dual quartet RBD-eVLPs) were lower against SARS-2 variants than against zoonotic sarbecoviruses (Fig. 2 C, Fig. S2 B). However, we recently reported increased neutralizing titers against hard-to-neutralize viruses including Omicron elicited by pulsatile release of a single injection of mosaic-8 RBD-NPs using atomic layer deposition (ALD) compared with bolus prime and boost injections of the same immunogen (Cohen et al., 2025). ALD is also an effective delivery mechanism for mRNA vaccines (Coleman et al., 2025), offering the possibility that increased neutralizing potencies would be observed for ALD formulations of mRNA-based mosaic-8 immunogens. In addition, improved neutralization of SARS-2 variants might be achieved by changing the RBD composition of mosaic-8 immunogens; e.g., a computationally-designed mosaic-7 RBD-NP elicited stronger neutralizing and binding Ab responses than original mosaic-8 RBD-NPs (Wang et al., 2025), and in independent studies of the immunogenicity of different mosaic RBD immunogen compositions, we observed increased cross-reactive responses when we removed the SARS-2 RBD from mosaic RBD immunogens (Cohen et al., 2024; Wang et al., 2025). Thus, altering the composition of dual RBD quartets (e.g., adding an Omicron RBD to an RBD quartet, adding clade 3 RBDs, and/or removing all SARS-2 RBDs) might improve neutralization potencies against SARS-2 variants.

In the absence of neutralization, binding Ab responses can trigger immune responses to clear infected cells through Fc-mediated effector functions (Paciello et al., 2024) and/or complement activation (Abu-Humaidan et al., 2022), and binding Ab titers correlate with protection in humans and animals vaccinated with COVID-19 mRNA vaccines (Corbett et al., 2021; Gilbert et al., 2022; Goldblatt et al., 2022). Indeed, human mAbs that neutralized early SARS-2 strains but lost neutralizing activity against SARS-2 Omicron variants retained Fc effector functions (Paciello et al., 2024), and non-neutralizing human anti-RBD mAbs protected from a SARS-2 challenge in animal models (Clark et al., 2024). In addition, T cell responses conferred protection from an Omicron challenge in the absence of neutralizing Ab responses (Case et al., 2024), and CD4+ T cells producing IFN-γ play critical roles in controlling SARS-CoV-2 infection independently of Abs (Fumagalli et al., 2024). Thus, even in the absence of strong neutralizing Ab titers against SARS-2 variants, the ability of mosaic-8 immunogens to elicit Abs that exhibit broad binding across sarbecovirus RBDs and balanced T cell responses are desirable traits for a protective pan-sarbecovirus vaccine.

For characterizing dual quartet immunogens, we describe a new systems serology technique, SySPEM, that quantifies the degree of binding to individual epitopes on an antigen across IgG subclasses and FcγR-binding IgGs, therefore allowing comparisons of epitope profiles elicited by different immunogens that are not possible using traditional systems serology (Ackerman et al., 2017). SySPEM allows higher throughput epitope mapping than, e.g., EMPEM (Turner et al., 2023), but can deliver some of the same information while also delineating epitopes recognized by different Ab classes. In contrast to epitope mapping competition ELISAs (Morris, 1996), multiplexed SySPEM analyses use minimal sample volumes, do not rely on a mAb panel of known epitopes, and provide more precise epitope definitions based on locations of introduced N-glycans. SySPEM and DMS results are complementary, but SySPEM experiments are easier to execute, require less material, and can allow quantitative comparisons between responses to distinct epitopes for different immunogens and IgG classes. Also, the SySPEM results reported here yielded mechanistic insights that were not revealed by DMS (e.g., quantifying differences in responses to dual quartet RBD-mRNA and dual quartet RBD-EABR-mRNA). In addition, we used SySPEM to show that the same antigen (dual quartet RBDs) delivered through mRNA- or protein-based vaccine modalities elicited distinct IgG class-specific epitope profiles, suggesting that the way an antigen is presented can impact its epitope recognition by the immune system.

SySPEM revealed potential advantages of the mRNA-EABR approach compared with a traditional mRNA immunogen; e.g., although overall binding and neutralizing Ab titers were similar, the dual quartet RBD-EABR-mRNA immunogen elicited stronger targeting of conserved class 4 and class 5 epitopes and weaker targeting of variable class 1/class 2 epitopes than dual quartet RBD-mRNA, suggesting that addition of an EABR sequence could improve the effectiveness of an mRNA-encoded dual quartet RBD pan-sarbecovirus vaccine. For example, immunogen presentation on circulating eVLPs would promote wider antigen distribution to potentially engage more B cells (Fan et al., 2025a; Hoffmann et al., 2023). Indeed, circulating RBD-eVLPs resulting from a dual quartet RBD-EABR-mRNA immunogen offer similar advantages to a recently described mRNA-delivered RBD protein nanoparticle (Hendricks et al., 2025) but also encode membrane-bound antigens at cell surfaces that could trigger additional immune responses. eVLP budding in the EABR approach might facilitate continuous replenishment of the cell surface with antigen, potentially increasing overall levels of antigen being displayed. The small size of eVLPs compared with a cell also facilitates efficient internalization by antigen-presenting cells (Foged et al., 2005; Joshi et al., 2013; Kim et al., 2022; Zepeda-Cervantes et al., 2020), which could result in enhanced antigenic peptide presentation by class II MHC and activation of T follicular helper cells to promote B cell activation and germinal center formation (Fan et al., 2025a). Improvements for mRNA vaccines by including EABR appear to be general, as the EABR approach enhances vaccine-induced humoral responses against SARS-2 (Cohen et al., 2021a; Cohen et al., 2024; Cohen et al., 2022; Hills et al., 2024; Wang et al., 2025), RSV (Chai et al., 2025), and influenza (Zhang et al., 2025). We also showed that a bivalent Wuhan-Hu-1/BA.5 spike-EABR-mRNA booster elicited more balanced targeting of multiple RBD epitopes in pre-vaccinated mice than conventional monovalent and bivalent spike-mRNA boosters, which primarily targeted variable RBD epitopes (Fan et al., 2025a). Taken together, our findings suggest that EABR-mRNA delivery of mosaic and multivalent immunogens improves targeting of conserved epitopes to enhance Ab breadth.

Pan-sarbecovirus vaccine candidates evaluated in mice would ideally elicit broad-binding IgG2a and IgG2b subclasses that promote robust Fc effector functions (Bruhns and Jonsson, 2015; Crescioli et al., 2025) to target conserved epitopes and elicit IgGs that bind to activating Fc receptors (FcγR3 and FcγR4). Using SySPEM, we showed that dual RBD quartet immunogens, including the two mRNA-encoded versions, fulfill these criteria. There are many future potential applications for this new technology for design and evaluation of vaccines against a variety of viruses and other pathogens. For example, studies to establish correlates of protection of vaccines could include SySPEM evaluations to determine relative protective effects of epitope targeting by different classes of IgGs, thus establishing desirable parameters for future vaccine efforts. In addition, SySPEM can be used to determine whether some epitopes elicit IgGs with different potential Fc effector functions to inform immunogen design for eliciting optimal immune responses against a particular pathogen.

In summary, our data support advancement of dual quartet RBD-mRNA immunogens with an optimal RBD composition into clinical development as a next-generation booster vaccine to provide protection from emerging SARS-2 variants and to reduce the chances of a future zoonotic sarbecovirus spillover from causing a new epidemic or pandemic.

## Methods

### Membrane-bound RBD quartet constructs

Sequences of the extracellular domains of RBD quartets 4a and 4b were based on previously reported SpyTag-Quartet (Genbank PP13603) and SpyTag-Alternate Quartet (GenBank PP136032, Addgene plasmid ID 214728) sequences (Hills et al., 2024). mRNA constructs encoded a mouse Ig heavy chain signal peptide, codon-optimized RBD quartets 4a and 4b in which individual RBDs were separated by nine-residue Gly/Ser/Thr linkers followed by a five-residue Gly/Ser linker, the SARS-1 spike transmembrane domain, and a truncated spike cytoplasmic tail (McBride et al., 2007) (GenBank AAP13441.1, residues [1218-1235]). For RBD quartet 4a-EABR and 4b-EABR constructs, an endocytosis-preventing motif (EPM) (Hoffmann et al., 2023), four-residue Gly/Ser linker, and the EABR (Hoffmann et al., 2023) sequence were appended to the C-terminus. For in vitro assays, these constructs were cloned into the IVTpro T7 plasmid (Takara 6144), containing the immunogen open reading frame flanked by 5′ and 3′ untranslated regions (UTRs) and a genetically encoded poly(A) tail.

### mRNA synthesis

For in vitro assays, IVTpro T7 plasmids encoding codon-optimized RBD quartet 4a, RBD quartet 4b, RBD quartet 4a-EABR, and RBD quartet 4b-EABR constructs were linearized by BspQI restriction digestion (NEB, R0712) and subsequently purified using the Zymo DNA Clean & Concentrator-25 kit (Zymo, D4034). mRNA was generated by in vitro transcription using the IVTpro™ T7 mRNA Synthesis Kit (Takara, 6144) following manufacturer’s instructions, with the addition of the CleanCap Reagent AG (3’ OMe) (TriLink, N-7413) and complete substitution of uridine with N1-Methylpseudouridine-5’-Triphosphate (TriLink, N-108; TriLink) to reduce immunogenicity (Hoffmann et al., 2023). mRNA was then purified by lithium chloride precipitation and an ethanol wash following standard protocols (Walker and Lorsch, 2013). Purified mRNA was resuspended in nuclease-free water and stored at –80 °C.

For mRNAs used for in vivo immunization studies, constructs were synthesized by RNAcore (https://www.houstonmethodist.org/research-cores/rnacore/) using proprietary manufacturing protocols including incorporation of the CleanCap Reagent AG (3’ OMe) (TriLink, N-7413) and complete substitution of uridine with N1-Methylpseudouridine-5’-Triphosphate (TriLink, N-108). mRNAs were purified by oligo-dT affinity purification. To remove double-stranded RNA contaminants, mRNAs were further purified using a cellulose-based method as described (Baiersdorfer et al., 2019). In brief, cellulose fibers (0.2 g/mL) were equilibrated in chromatography buffer (10mM HEPES pH 7.2, 0.1 mM EDTA, 125 mM NaCl, 16% ethanol) and added to microcentrifuge spin columns. mRNA diluted in chromatography buffer was added and shaken vigorously at room temperature for 30 minutes. The flow-through containing single-stranded mRNA was collected and purified with a sodium acetate precipitation and an ethanol wash. The final purified mRNA was filtered through a 0.2 µm filter before storage at –80 °C (Baiersdorfer et al., 2019; Hoffmann et al., 2023).

### mRNA transfections and flow cytometry

HEK293T cells were seeded at 1 × 10⁶ cells per well in 6-well plates and incubated for 20 h before transfection with 2 µg of mRNA encoding RBD quartet 4a, RBD quartet 4b, and/or their EABR-fused counterparts (4a-EABR, 4b-EABR) using Lipofectamine™ MessengerMax™ (Thermo Fisher Scientific). Forty-eight hours post-transfection, supernatants were collected for eVLP purification by ultracentrifugation on a 20% sucrose cushion as described (Hoffmann et al., 2023). Cells were gently detached, resuspended in PBS+ (PBS + 2 % FBS), and 100 µL aliquots were taken for flow cytometry.

For the experiment shown in Fig. S1 A, cells were stained with the cross-reactive anti-RBD mAb C118 (Jette et al., 2021; Robbiani et al., 2020) (2.5 µg/mL) for 30 min in the dark at room temperature, washed, and subsequently stained with Alexa Fluor® 647-conjugated anti-human IgG secondary Ab (Invitrogen, A21445; 1:4,000) for 15 min. After a final wash, cells were resuspended in PBS+ and analyzed on an Attune NxT Flow Cytometer (Thermo Fisher).

For the experiment shown in Fig. S1 C, cells were either singly transfected or co-transfected with combinations of RBD quartet 4a and 4b constructs and stained simultaneously with 2.5 µg/mL of Alexa Fluor® 647-conjugated anti-Rs4081 RBD (Fan et al., 2025b) and Alexa Fluor® 488-conjugated M8a-7 IgG (anti-WIV1 RBD) (Fan et al., 2022) mAbs for 30 min at room temperature in the dark. After washing and resuspension, samples were analyzed on an Attune NxT Flow Cytometer. Results were plotted using FlowJo 10.5.3 software.

### Production of dual quartet and SARS-2 RBD-eVLPs

Dual quartet RBD-eVLP and SARS-2 RBD-eVLP samples were generated as described (Hoffmann et al., 2023) by transfecting Expi293F cells (Gibco) cultured in Expi293F expression media (Gibco) on an orbital shaker at 37°C and 8% CO_2_. To generate dual quartet RBD-eVLPs, DNA plasmids encoding RBD quartet 4a-EABR and RBD quartet 4b-EABR constructs were mixed at a ratio of 1:1. 72 hours post-transfection, cells were centrifuged at 400 x g for 10 min, supernatants were passed through a 0.45 μm syringe filter and concentrated using Amicon Ultra-15 centrifugal filters with 100 kDa molecular weight cut-off (Millipore). eVLPs were purified by ultracentrifugation at 50,000 rpm (135,000 x g) for 2 hours at 4°C using a TLA100.3 rotor and an Optima^TM^ TLX ultracentrifuge (Beckman Coulter) on a 20% w/v sucrose cushion. After removal of supernatants by aspiration, pellets were re-suspended in 200 μL sterile PBS at 4°C overnight. To remove residual debris, samples were centrifuged at 10,000 x g for 10 min, and supernatants were collected. eVLPs were further purified by SEC using a Superose 6 Increase 10/300 column (Cytiva) equilibrated with PBS. Peak fractions corresponding to eVLPs were combined and concentrated to 250-500 μL in Amicon Ultra-4 centrifugal filters with 100 kDa molecular weight cut-off. Samples were aliquoted and stored at -20°C.

The amount of SARS-2 RBD on SARS-2 RBD-eVLPs and the amount of dual quartet RBD on dual quartet RBD-eVLPs were determined by quantitative Western blot analysis. Various dilutions of SEC-purified eVLP samples and known amounts of either soluble SARS-2 RBD protein (Sino Biological, 40591-V08H-B-20) or purified soluble SpyTagged dual quartet RBDs (SpyTagged RBD quartets 4a and 4b were mixed at a ratio of 1:1) were separated by SDS-PAGE and transferred to nitrocellulose membranes (0.45 μm) (Thermo Fisher Scientific, LC2001). Rabbit anti-SARS-2 S1 polyclonal IgG (Thermo Fisher Scientific, PA5-116916; 1:1,000) and HRP-conjugated goat anti-rabbit IgG (Abcam, ab98467; 1:10,000) were used. Protein bands were visualized using ECL Prime Western Blotting Detection Reagent (Cytiva, RPN2232). Band intensities of the respective standards and eVLP sample dilutions were measured using ImageJ to determine RBD amounts.

### LNP encapsulation of mRNAs

Purified dual quartet RBD-encoding mRNAs (RBD quartet 4a + RBD quartet 4b or RBD quartet 4a-EABR + RBD quartet 4b-EABR; 1:1 ratio) for in vivo studies were formulated in LNPs as previously described (Pardi et al., 2015). 1,2-distearoyl-*sn*-glycero-3-phosphocholine, cholesterol, a PEG lipid, and an ionizable cationic lipid dissolved in ethanol were rapidly mixed with an aqueous acidic solution containing mRNA using an in-line mixer. The ionizable lipid and LNP composition are described in the international patent application WO2017075531(2017). The post in-line solution was dialyzed with PBS to remove the ethanol and displace the acidic solution. LNP was measured for size (60-65 nm) and polydispersity (PDI < 0.075) by dynamic light scattering (Malvern Nano ZS Zetasizer). Encapsulation efficiencies were >97% as measured by the Quant-iT Ribogreen Assay (Invitrogen).

### Protein expression

RBDs, RBD quartets, and soluble sarbecovirus spike-6P (Hsieh et al., 2020) (SARS-2 WA1 and JN.1) or spike-2P (XBB.1.5, SARS-1, Rf1, Rs4091, Yun11, BM48-31, BtKY72) trimers were expressed as described (Cohen et al., 2025; Cohen et al., 2022; Hills et al., 2024). Briefly, AviTag and/or His-tagged proteins were purified from transiently-transfected Expi293F cells (Gibco) by nickel affinity chromatography (HisTrap HP, Cytiva) and SEC (Superose 6 Increase 10/300, Cytiva) (Barnes et al., 2020; Cohen et al., 2022; Wang et al., 2022). Fractions corresponding to proteins of interest were pooled, concentrated, and stored at 4°C. Biotinylated proteins were generated as described (Tykvart et al., 2012) by co-transfection of AviTag/His-tagged spike and RBD constructs with a plasmid encoding an endoplasmic reticulum-directed BirA enzyme (kind gift from Michael Anaya, Caltech). The anti-Rs4081, anti-RmYN02, and anti-WIV1 RBD mAb IgGs from Fig. S1 and the mAb IgGs and human ACE2-Fc protein shown in Fig. S7 C were expressed, purified, and classified as recognizing the RBD epitopes as described (Barnes et al., 2020; Fan et al., 2022; Fan et al., 2025b; Jette et al., 2021).

### Preparation of RBD-NPs

For making dual quartet RBD-NPs, SpyCatcher003-mi3–C-Tag (Addgene 159995) subunits were expressed in BL21 (DE3) *E. coli* (Agilent) as described (Rahikainen et al., 2021). SpyCatcher003-mi3–C-Tag bacterial cell pellets were resuspended in 20 mL of 20 mM Tris-HCl pH 8.5, 300 mM NaCl, supplemented with 0.1 mg/mL lysozyme, 1 mg/mL cOmplete mini EDTA-free protease inhibitor (Roche), and 1.0 mM PMSF. After incubation at 4 °C for 45 min with end-over-end mixing, an Ultrasonic Processor equipped with a microtip (Cole-Parmer) was used to perform sonication on ice (four times for 60 s, 50% duty-cycle), and cell debris were cleared by centrifugation (35,000 x g for 45 min at 4 °C). SpyCatcher003-mi3–C-Tag was purified by ammonium sulfate precipitation (170 mg per mL of lysate) and resuspension, followed by SEC using a HiPrep Sephacryl S-400 HR 16-60 column (GE Healthcare) equilibrated with PBS as described (Rahikainen et al., 2021). Dual quartet RBD-NPs were generated by incubating purified SpyCatcher003-mi3–C-Tag with a 1:1 molar ratio of purified SpyTagged RBD quartets at 4 °C in PBS.

### Immunization of mice

Mouse procedures were approved by the Labcorp Institutional Animal Care and Use Committee. Six- to 7-week-old female BALB/c mice from Charles River Laboratories (RRID: IMSR_JAX:000664) were housed at Labcorp Drug Development, Denver, PA for immunizations. All animals were healthy upon receipt and monitored during a 7-day acclimation period. Mice were assigned randomly to experimental groups of 10 animals. Mouse cages were kept in a climate-controlled room (68 - 79 °C) at 50 ± 20% relative humidity and provided with Rodent Diet #5001 (Purina Lab Diet) ad libitum.

Mice were immunized by intramuscular injection on days 0 and 28 with 1.5 or 0.5 µg dual quartet RBD-mRNA, 1.5 or 0.5 µg dual quartet RBD-EABR-mRNA, 5 µg dual quartet RBD-NP, 5 µg dual quartet RBD-eVLP, or 5 µg SARS-2 RBD-eVLP (based on RBD content for protein-based immunogens). Prior to immunizations, dual quartet RBD-NP, dual quartet RBD-eVLP, and SARS-2 RBD-eVLP immunogens were mixed with Addavax adjuvant (InvivoGen; 50% v/v). Serum samples for ELISAs and neutralization assays were obtained on indicated days by retro-orbital bleeding.

### ELISAs

For eVLP sandwich ELISAs, 96-well high-binding plates (Corning, #9018) were coated overnight at 4 °C with capture mAb (5 µg/mL in 0.1 M NaHCO₃, pH 9.8). Plates were blocked with 3% BSA and 0.1% Tween-20 in TBS (TBS-T) for 30 min at room temperature, followed by addition of serially diluted purified eVLP samples for 2 h. After washing, biotinylated detection Abs (5 µg/mL in blocking buffer) were added for 2 h at room temperature. Plates were washed three times with TBS-T, incubated with streptavidin-HRP (Abcam, 7403; 1:20,000) for 30 min, washed again, and developed with 1-Step™ Ultra TMB-ELISA substrate solution (ThermoFisher). Reactions were stopped with 1 N HCl and absorbance was measured at 450 nm.

For the ELISA in Fig. S1 B, plates were coated with either anti-Rs4081 (Fan et al., 2025b) or anti-RmYN02 (Fan et al., 2025b) mAbs and detected with biotinylated C118 (Jette et al., 2021; Robbiani et al., 2020). For the ELISA in Fig. S1 D, plates were coated with anti-WIV1 (Fan et al., 2022) and detected with biotinylated anti-Rs4081 (Fan et al., 2025b).

For serum ELISAs, purified His-tagged RBD (2.5 μg/mL in 0.1 M sodium bicarbonate buffer pH 9.8) was coated onto Nunc® MaxiSorp™ 384-wellplates (Sigma) and incubated overnight at 4°C. After blocking with 3% bovine serum albumin (BSA), 0.1% Tween 20 in Tris-buffered saline (TBS-T) for 1 hour at room temperature, blocking solution was removed by aspiration. For serum ELISAs shown in Fig. 2 and Fig. S2, mouse serum (1:100 initial dilution) was serially diluted by 3.1-fold in TBS-T/3% BSA and then added to plates for 3 hours at room temperature, followed by washing with TBS-T. Plates were then incubated for 1 hour with a 1:100,000 dilution of HRP-conjugated anti-IgG secondary (goat anti-mouse IgG; Abcam; RRID: AB_955439), and plates were washed with TBS-T. For mAb ELISAs shown in Fig. S7, purified mAbs were serially diluted by 3.1-fold in TBS-T/3% BSA and then added to plates for 3 hours at room temperature, followed by washing with TBS-T. Plates were then incubated for 1 hour with a 1:100,000 dilution of HRP-conjugated anti-IgG secondary (goat anti-human IgG; SouthernBiotech 2014-05), and plates were washed with TBS-T. SuperSignal™ ELISA Femto Substrate (ThermoFisher) was added as per manufacturer’s instructions, and luminescence was read at 425 nm. Data were collected in duplicate and midpoint titers (ED_50_ values) were obtained using Graphpad Prism 10.5.0 assuming a four-parameter dose-response curve fit.

### Pseudovirus neutralization assays

Lentivirus-based pseudoviruses were produced in HEK293T cells (RRID:CVCL_0063) cultured in Dulbecco’s modified Eagle’s medium (DMEM, Gibco) supplemented with 10% heat-inactivated fetal bovine serum (FBS, Bio-Techne), 1% Penicillin/Streptomycin (Gibco), and 1% L-Glutamine (Gibco) at 37 °C and 5% CO_2_. HEK293T-hACE2 cells (RRID:CVCL_A7UK) (Starr et al., 2021b) and high-hACE2 HEK-293T (kindly provided by Kenneth Matreyek, Case Western Reserve University) target cells for neutralization assays were cultured in DMEM (Gibco) supplemented with 10% heat-inactivated FBS (Bio-Techne), 5 mg/mL gentamicin (Sigma-Aldrich), and 5 mg/mL blasticidin (Gibco) at 37 °C and 5% CO_2_.

Lentiviral-based viruses were prepared as described (Crawford et al., 2020; Robbiani et al., 2020) using genes encoding spike sequences lacking C-terminal residues in the cytoplasmic tail: deletions of 21 residues (SARS-2 variants) or 19 residues (SARS-1, Khosta-2-SARS-1 chimera, BtKY72-SARS-1 chimera). BtKY72 (containing K493Y/T498W substitutions) and Khosta-2 pseudoviruses were made with chimeric spikes in which the RBD from SARS-1 (residues 323-501) was substituted with the RBD from BtKY72 K493Y/T498W (residues 327-503) or Khosta-2 (residues 324-500) as described. (Seifert et al., 2022). Cells were co-transfected with HIV-1-based lentiviral plasmids, a luciferase reporter gene, and a coronavirus spike construct, resulting in lentivirus-based pseudovirions expressing a sarbecovirus spike protein. Supernatants were harvested 48-72 hours post-transfection, filtered, and stored at -80 °C. Pseudovirus infectivity was determined by titration using HEK293T-hACE2 cells.

Prior to neutralization assays, serum from immunized mice was heat-inactivated at 56 °C for 10 minutes. Heat-inactivated serum was three-fold serially diluted and incubated with pseudovirus for 1 hour at 37°C, then the serum/virus mixture was added to HEK293T-hACE2 target cells or high-hACE2 HEK-293T cell line expressing hACE2 encoded with a consensus Kozak sequence (for SHC014 assays; kindly provided by Kenneth Matreyek, Case Western Reserve University) and incubated for 48 hours at 37°C. After removing media, cells were lysed with Britelite Plus reagent (Revvity Health Sciences), and luciferase activity was measured as relative luminesce units (RLUs). Relative RLUs were normalized to RLUs from cells infected with pseudotyped virus in the absence of antiserum. Half-maximal inhibitory dilutions (ID_50_ values) were derived in AntibodyDatabase (West et al., 2013) using 4-parameter nonlinear regression.

### ELISpot assays

Mice were euthanized on day 77 or 78, and spleens were collected and processed as described (Hoffmann et al., 2023). Briefly, spleens were homogenized using a gentleMACS Octo Dissociator (Miltenyi Biotec). Cells were passed through a 70 µm tissue screen, centrifuged at 1,500 rpm for 10 min, and resuspended in CTL-Test^TM^ media (ImmunoSpot) containing 1% GlutaMAX^TM^ supplement (Gibco). ELISpot assays were performed as described (Hoffmann et al., 2023). A SARS-CoV-2 spike PepMix^TM^ pool of 315 peptides (15-mers with 11 amino acid overlap; JPT Peptide Technologies) was added to mouse IFNγ/IL4 double-color ELISpot plates (ImmunoSpot) at a concentration of 2 µg/mL, and 300,000 splenocytes were added per well. Plates were incubated at 37°C for 24 hours. Biotinylated detection reagents, streptavidin-alkaline phosphatase (AP), and substrate reagents were added according to the manufacturer’s guidelines. To stop the reactions, plates were gently rinsed with water three times. Plates were air-dried for two hours in a running laminar flow hood. Numbers of spots were quantified using a CTL ImmunoSpot S6 Universal-V Analyzer (Immunospot).

### DMS Epitope Mapping

DMS experiments to map epitopes recognized by serum Abs were performed in biological duplicates using independent mutant RBD yeast libraries (WA1 (Starr et al., 2022) and SARS-1 (Starr et al., 2022) generously provided by Tyler Starr, University of Utah) as described (Cohen et al., 2022; Greaney et al., 2021d). Sera that had been heat inactivated for 30 min at 56 °C were incubated twice with 50 OD units of AWY101 yeast transformed with an empty vector in order to remove non-specific yeast-binding Abs. Expression of RBDs in libraries was induced in galactose-containing synthetic defined medium with Casamino acids (6.7g/L Yeast Nitrogen Base, 5.0 g/L Casamino acids, 1.065 g/L MES acid, and 2% w/v galactose plus 0.1% w/v dextrose). After 18 hours of induction, cells were washed twice and incubated with serum under conditions of gentle agitation for 1 hour at room temperature. Cells were labeled for 1 hour with a secondary Ab (1:200 Alexa Fluor-647-conjugated goat anti-mouse-IgG Fc-gamma, Jackson ImmunoResearch 115-605-008, RRID:AB_2338904) after washing twice.

Stained yeast cells were examined by fluorescence-activated cell sorting (FACS) using a Sony SH800 cell sorter. Cells were gated to capture RBD mutants that showed reduced Ab binding compared to a control. Cells were collected for each sample until ∼5 x 10^6^ RBD^+^ cells were processed, which corresponds to ∼5 x 10^5^- 1 x 10^6^ RBD^+^ Ab-escaped cells. These cells were grown overnight in synthetic defined media (6.7 g/L Yeast Nitrogen Base, 5.0 g/L Casamino acids, 1.065 g/L MES acid, and 2% w/v dextrose, 100 U/mL penicillin, 100 µg/mL streptomycin) to expand cells prior to plasmid extraction. DNA extraction and Illumina sequencing were done as described (Hills et al., 2024) (raw sequencing data will be available on NCBI SRA). Escape fractions were computed as described (Greaney et al., 2021d; Hills et al., 2024) using Swift DMS (Hills et al., 2024) (available upon request). Escape scores were calculated using a filter to remove variants with mutations that escaped binding because of poor expression, >1 amino acid mutation, or low sequencing counts (Greaney et al., 2022; Hills et al., 2024).

Escape map visualizations (static line plots, logo plots, and structural depictions) were created using Swift DMS (Hills et al., 2024). Line heights show the escape score for a particular amino acid substitution, as described (Hills et al., 2024). In some visualizations, sites were categorized based on RBD epitope region (Barnes et al., 2020; Cui et al., 2024; Jensen et al., 2023; Jette et al., 2021): class 1 (pink; RBD residues 403, 405, 406, 417, 420, 421, 453, 455-460, 473-478, 486, 487, 489, 503, 504); class 2 (purple; residues 472, 479, 483-485, 490-495), class 3 (blue; residues 341, 345, 346, 437-450, 496, 498-501,), class 4 (orange; residues 365-390, 408), class 5 (green; residues 352-357, 396, 462-468). In structural depictions of DMS results, an RBD surface (PDB 6M0J) was colored by the site-wise escape metric at each site, with red scaled at an escape fraction of 2.0. Residues exhibiting the highest escape fractions were labeled with their residue number, which was colored according to epitope class. Logo plot residues are colored according to RBD epitopes within different classes as indicated on the legend.

We stratified DMS escape fraction values into four groups. Escape fractions for each RBD substitution range from 0 (no cells with this substitution were sorted into the escape bin) to 1 (all cells with this substitution were sorted into the serum Ab escape bin) (Greaney et al., 2021c). The sum of escape fractions for all substitutions at a specific site is represented by the total escape peak (Greaney et al., 2021c). DMS profiles with total escape peaks <0.5 at all sites were classified as polyclass responses. DMS profiles with total escape peaks of 0.5 to 1, >1 to 2, or >2 at one or more sites were classified as weak, moderate, or strong escape profiles, respectively, corresponding to their RBD epitope (Table 1).

### Systems serology: Ab subclass and FcγR binding profiling

Serum samples from immunized mice were analyzed in systems serology (Ackerman et al., 2017) experiments as described (Cohen et al., 2025) using modifications of a Luminex assay to quantify the levels of antigen-specific Ab subclasses and FcγR binding profiles, which allows for simultaneous detection and quantification of multiple analytes (i.e., different antigens) in a single sample (Brown et al., 2017). In brief, avidin (Sigma-Aldrich Catalog #: A9275-25MG) was coupled by carbodiimide-NHS ester chemistry to magnetic Luminex microspheres (LuminexCorp) using Sulfo-NHS (ThermoFisher™ Catalog number 24510) and 1-Ethyl-3-[3-dimethylaminopropyl]carbodiimide hydrochloride (EDC) (ThermoFisher™ Catalog number 22980) according to the manufacturer’s instructions. Microspheres were blocked for 30 min with 1% BSA in PBS pH 7.4 (assay buffer) and then washed twice with PBS. Microspheres were incubated with biotinylated antigen (sarbecovirus spike trimer, RBD, or control antigen) at room temperature for 2 hours in assay buffer followed by blocking with 10 µM biotin (Millipore Sigma, Catalog B4501-100MG). Antigen-coupled microspheres were combined then incubated with heat-inactivated serum samples for 1 hour at room temperature in 96-well plates (Corning) at 1:200 or 1:1000 dilutions. Unbound IgGs were removed by washing twice with 200 µL of assay buffer. Total IgG was detected using a PE-conjugated secondary Ab recognizing mouse IgG (Southern Biotech 1030-09S). For evaluating antigen binding by different IgG subclasses, secondary Abs against mouse IgG subclasses (Southern Biotech 1070-09S, 1080-09S, 1090-09S, 1100-09S; PE-coupled anti-IgG1, IgG2a, IgG2b, IgG3) were added at a 1:1000 dilution in assay buffer and incubated for 1 hour with continuous shaking at room temperature. Excess primary and secondary Abs were removed by washing twice with 200 µL of assay buffer. Beads were resuspended in 200 µL of assay buffer, run in duplicate, and flowed in single file past two lasers on a Luminex™ FLEXMAP 3D™ Instrument System. In a Luminex assay, each bead region is impregnated with a different ratio of fluorescent dyes such that bead region identity, and thereby antigen identity, is interrogated by the first laser. A second laser interrogates the signal of secondary Ab binding. Median fluorescence intensity (MFI) was calculated for all samples.

For evaluating FcγR-binding IgGs, soluble ectodomains of 6xHis-tagged FcγR2b, FcγR3, and FcγR4 were prepared as described (Cohen et al., 2025) and biotinylated by co-expression with BirA enzyme in Expi293T cells (Tykvart et al., 2012). Biotinylated FcγRs were purified on a HisTrap column (VWR) according to the manufacturer’s instructions followed by SEC and then bound to PE-streptavidin (eBioscience). Labeled FcγRs were diluted 1:200 in assay buffer and incubated with serum-coated microspheres for 1 hour at room temperature with continuous shaking. Unbound primary and PE-labeled FcγR were removed as described for anti-IgG subclass secondary Abs. Beads were resuspended and run on a Luminex™ FLEXMAP 3D™ Instrument System to derive MFI values as described above.

Log₁₀-transformed heatmaps of antigen-specific IgG responses were generated using Python (v3.9.16) with the Pandas (v2.0.3), NumPy (v1.23.5), Matplotlib (v3.7.1), and Seaborn (v0.12.2) packages. A soluble influenza hemagglutinin trimer (listed as CA-09 HA in Fig. 5; expressed and purified as described (Cohen et al., 2021b)) was used as a control to evaluate non-specific binding.

### SySPEM

One or more potential N-linked glycosylation site sequons (Asn-x-Ser/Thr) (Lehle and Bause, 1984) were introduced at solvent-exposed residue(s) within the class 1 and class 2, class 2, class 3, class 4, class 1/4, or class 5 RBD epitopes (Barnes et al., 2020; Cui et al., 2024; Jensen et al., 2023; Jette et al., 2021) of the WA1 RBD (Fig. S7 A). Addition of N-glycan(s) was confirmed by SDS-PAGE as a shift in electrophoretic mobility to a higher apparent molecular weight (Fig. S7 B). Proper folding of RBD KO mutants and expected effects of added N-glycans were verified by assaying the binding of characterized mAbs or a human ACE2-Fc construct (Jette et al., 2021) by ELISA (Fig. S7 C). Binding of serum subsets to RBD wt and RBD KO proteins was measured using a Luminex multiplexed bead-based immunoassay as described above. The SySPEM score for each IgG sample/epitope pair was calculated as [1.0 – (RBD KO binding / RBD wt binding) x 100]. SySPEM scores range from limits of 100 (none of the IgGs in the sample bound to the RBD KO; i.e., all of the IgGs recognize the epitope that was targeted by the glycan addition) to 0 (all of the IgGs in the sample bound to the RBD KO; i.e., none of the IgGs recognize the epitope that was targeted by the glycan addition) (Fig. S6). Python (v3.9.16) with the Pandas (v2.0.3), NumPy (v1.23.5), Matplotlib (v3.7.1), and Seaborn (v0.12.2) packages were used for box and whisker plots of SySPEM data. For UMAP visualizations in Fig. 7, SySPEM scores of each serum sample were organized into a feature matrix where each column contains an epitope class SySPEM score, and each row corresponds to an individual sample. Features were standardized using the standard scalar function on sckit-learn 1.4.2. UMAP was then performed with umap-learn (0.5.9.post2) on the standardized matrix. UMAP parameters: the nearest neighbors (n-neighbors) parameter was set to 12 and minimum distance (min-dist) was set to 0.25 to preserve the best local and global structures on the UMAP plot. Seaborn (v0.12.2) was used to plot resulting UMAPs.

### Quantification and statistical analyses

We used pairwise comparisons, a method to evaluate sets of mean binding titers against individual viral strains for different immunization cohorts, as described previously (Cohen et al., 2024) to determine if results from different immunized cohorts were significantly different from each other. For ELISAs and neutralizations (Fig. 2), we used analysis of variance (ANOVA) followed by Tukey’s multiple comparison post hoc tests with the Geisser-Greenhouse correction, with pairing by viral strain, of ED_50_s/ID_50_s (converted to log_10_ scale) calculated using GraphPad Prism 10.5.0 to identify statistically significant titer differences between immunized groups for ELISAs. For comparisons of neutralizing titers by strain (Fig. S2), statistically significant titer differences between immunized groups for each given strain were determined using unpaired t test calculated using GraphPad Prism 10.5.0. For ELISpot assays, statistically significant differences between immunized cohorts were calculated using ordinary one-way ANOVA followed by Tukey’s multiple comparison test using GraphPad Prism.

For statistical analysis of systems serology results, responses were aggregated by computing the geometric mean (geomean) across replicate samples for each immunogen-antigen combination. Log₁₀-transformed geomeans were compared across immunogen groups using Tukey’s multiple comparison post hoc tests with the Geisser-Greenhouse correction, with pairing by viral strain calculated using GraphPad Prism 10.5.0. Box-and-whisker plots were generated for each IgG subclass or FcγR-binding IgG, displaying individual antigen-level geomeans per immunogen. Python (v3.9.16) with statsmodels (v0.14.4) packages were used for SySPEM Tukey’s HSD posthoc statistical analyses. Statistical significance was annotated using brackets and asterisk notation (p < 0.05 *, p < 0.01 **, p < 0.001 ***, p < 0.0001 ****).

### Materials, data, and code availability

DMS raw sequencing data is available on the NCBI SRA under BioProject PRJNA1067836, BioSample SAMN52937385. SySPEM scores for each IgG sample/epitope pair will be made publicly available upon publication. Materials are available upon request to corresponding authors with a signed material transfer agreement, and other information required to analyze the data in this paper is available from the lead contacts upon request. Code used for data processing and visualization of DMS, systems serology, and SySPEM results is available upon request. The work is licensed under a Creative Commons Attribution 4.0 International (CC BY 4.0) license, which permits unrestricted use, distribution, and reproduction in any medium, provided the original work is properly cited. To view a copy of this license, visit https://creativecommons.org/licenses/by/4.0/. This license does not apply to figures/photos/artwork or other content included in the article that is credited to a third party; obtain authorization from the rights holder before using such material.

## Acknowledgements

We thank Annie Rorick, Han Gao, and Priyanthi Gnanapragasam for experimental assistance, Jesse Bloom (Fred Hutchinson Cancer Research Center) and Tyler Starr (University of Utah) for RBD libraries, Ryan McNamara and Galit Alter (Ragon Institute) for help with Systems Serology experiments, Jost Vielmetter, Luisa Segovia, Alyssa Player, Annie Lam, and the Caltech Beckman Institute Protein Expression Center for protein production, Igor Antoshechkin and the Caltech Millard and Muriel Jacobs Genetics and Genomics Laboratory for Illumina sequencing, Michael Anaya for help with Luminex experiments, Dom Aylard, Katie Lippert, and the Gladstone Flow Cytometry Core for help with flow cytometry, and Anthony West and Ying Tam (Acuitas) for critical reading of the manuscript.

## Funding Declaration

These studies were funded by Wellcome Leap (P.J.B.), the National Institutes of Health P01-AI165075 (P.J.B.), DP5OD033362 (M.A.G.H.), and 1K99AI185267 (A.A.C.), the Medical Research Council (MRC grant no. MR/Y011910/1 to M.R.H.), the Merkin Institute for Translational Research (Caltech) (P.J.B.), and the James B. Pendleton Charitable Trust that supports the Gladstone Flow Cytometry Core. This manuscript is the result of funding in whole or in part by the National Institutes of Health (NIH). It is subject to the NIH Public Access Policy. Through acceptance of this federal funding, NIH has been given a right to make this manuscript publicly available in PubMed Central upon the Official Date of Publication, as defined by NIH.

## Author Contributions

Conceptualization: A.A.C., J.R.K., R.A.H., M.R.H., M.A.G.H., P.J.B.; Methodology: A.A.C., M.A.G.H.; Software: A.A.C.; Investigation: A.A.C., J.R.K., LM, IM, A.-C.P.F., K.-M.A.D., HES, M.A.G.H.; Resources: R.A.H., W.J.M., P.J.C.L.; Writing – original draft: A.A.C., J.R.K., K.-M.A.D., M.A.G.H., P.J.B.; Writing – review and editing: A.A.C., J.R.K., K.-M.A.D., R.A.H., M.R.H., M.A.G.H., P.J.B.; Visualization: A.A.C., J.R.K., K.-M.A.D., M.A.G.H., P.J.B.; Supervision: M.R.H., M.A.G.H., P.J.B.; Funding: M.A.G.H., P.J.B..

## Competing Interests

P.J.B. and A.A.C. are inventors on a US patent application (17/523,813) filed by the California Institute of Technology that covers mosaic RBD-NPs. M.A.G.H. and P.J.B. are inventors on a patent application (US17/835,751; international: PCT/US2022/032702) filed by the California Institute of Technology that covers the EABR technology. M.R.H. is an inventor on patents on spontaneous amide bond formation (EP2534484 and UK Intellectual Property Office 1706430.4) and a SpyBiotech co-founder and shareholder. P.J.B., M.A.G.H., A.A.C., J.R.K., K-M.A.D., R.A.H., and M.R.H. are co-inventors on a provisional US patent application filed by the California Institute of Technology that covers dual quartet mRNA immunogens. P.J.B. is a scientific advisor for Vaccine Company, Inc. M.A.G.H. is a scientific consultant for Vaccine Company, Inc. W.J.M. and P.J.C.L. are employees of Acuitas Therapeutics, a company developing LNP delivery technology; P.J.C.L. holds equity in Acuitas Therapeutics.

## Supplementary Information

**Figure S1.**
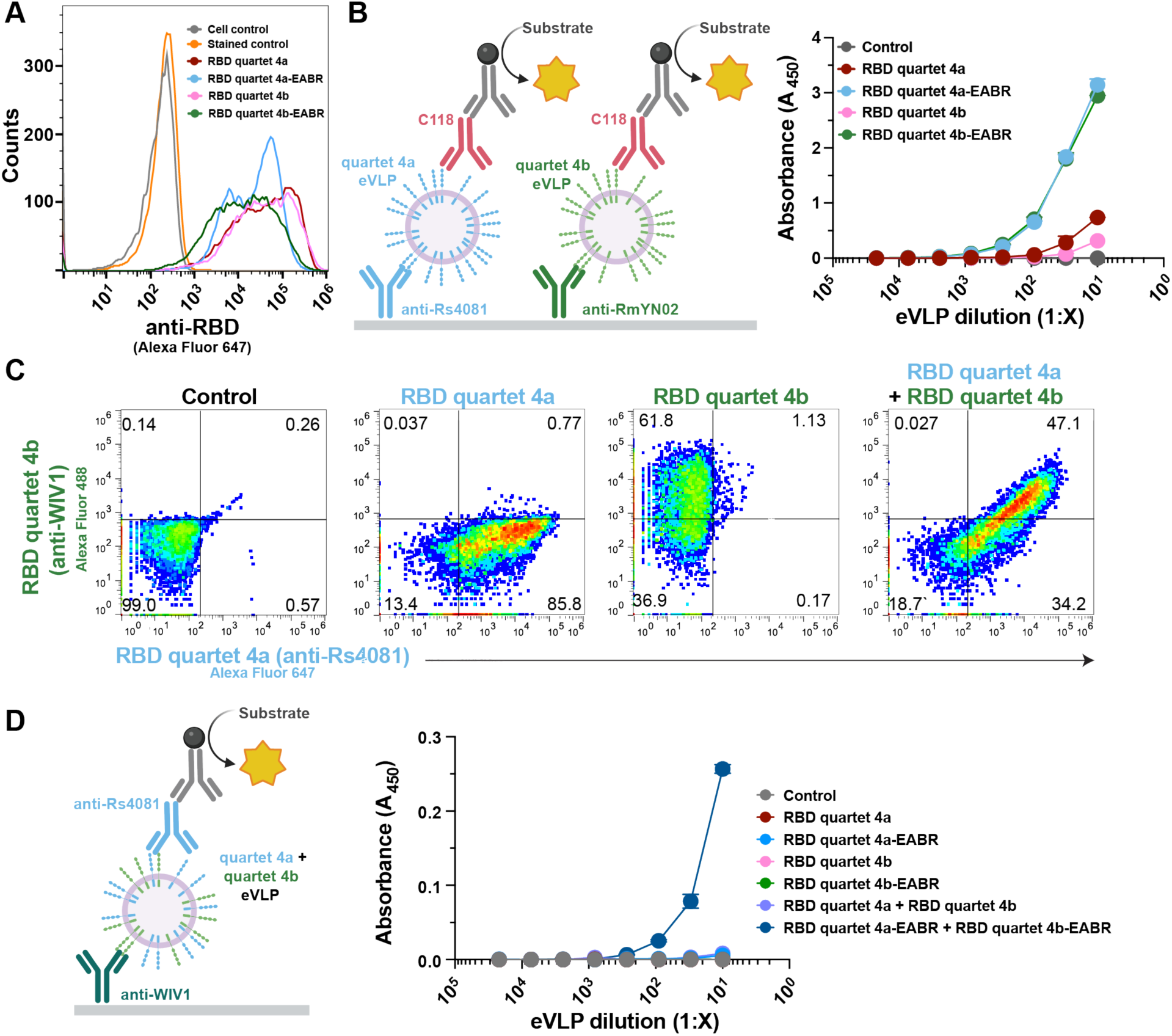
mRNA-encoded dual quartets are presented on cells and eVLPs. (A) Flow cytometry using C118, a human pan-sarbecovirus RBD-specific mAb (Jette et al., 2021; Robbiani et al., 2020), demonstrating that transfections of mRNAs encoding EABR and non-EABR versions of RBD quartet 4a or quartet 4b resulted in expression on the surface of transfected cells. Untransfected cells are shown as controls (unstained or stained with C118). (B) Left: ELISA schematic. RBD quartet levels were measured by capturing purified eVLPs with an anti-Rs4081 RBD mAb (Fan et al., 2025b) (recognizes RBD quartet 4a but not quartet 4b) or an anti-RmYN02 RBD mAb (Fan et al., 2025b) (recognizes RBD quartet 4b but not quartet 4a) and detecting with the pan-sarbecovirus RBD-specific C118 mAb (Jette et al., 2021; Robbiani et al., 2020). Right: ELISA showing that purified single quartet eVLPs purified from supernatants of cells transfected with mRNA encoding RBD quartet 4a-EABR or RBD quartet 4b-EABR displayed RBD quartets on their surface. The mean absorbance of two replicates is shown as a function of eVLP dilution with error bars representing standard deviations. (C) Flow cytometry demonstrating that RBD quartets 4a and 4b were co-expressed on the surfaces of individual cells. RBD quartet 4a was detected using an anti-Rs4081 RBD mAb (Fan et al., 2025b) and RBD quartet 4b was detected using anti-WIV1 RBD mAb (Fan et al., 2022). Untransfected cells stained with the anti-Rs4081 and anti-WIV1 RBD mAbs are shown as controls. The percentage of cells in each quadrant is shown. (D) Left: ELISA schematic. Purified eVLPs were evaluated in a sandwich ELISA in which an anti-WIV1 RBD mAb (Fan et al., 2022) (recognizes RBD quartet 4b but not quartet 4a) was used as a capture Ab and an anti-Rs4081 RBD mAb (Fan et al., 2025b) (recognizes quartet 4a but not quartet 4b) was used for detection. Right: ELISA showing detection of both RBD quartets 4a and 4b on the surfaces of eVLPs purified from supernatants of cells co-transfected with mRNAs encoding RBD quartet 4a-EABR and 4b-EABR constructs. The mean absorbance of two replicates is shown as a function of eVLP dilution with error bars representing standard deviations. Binding was not detected for purified supernatant samples from untransfected cells (control) or from supernatants from cells transfected with RBD quartet 4a, RBD quartet 4a-EABR, RBD quartet 4b, RBD quartet 4b-EABR, or RBD quartet 4a plus RBD quartet 4b (all data were plotted but some data points near 0.0 A_450_ are obscured by others).

**Figure S2.**
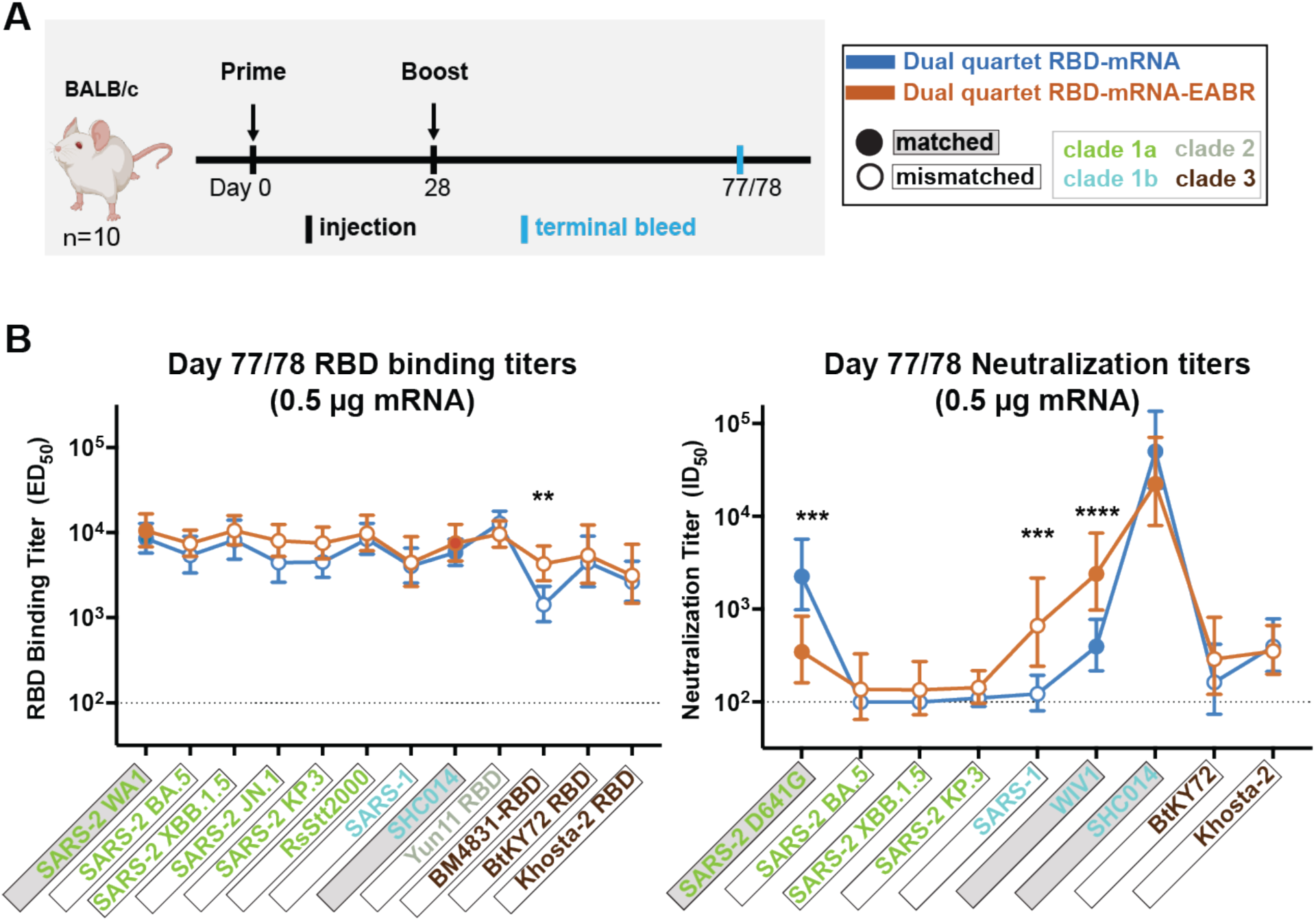
Dual quartet mRNAs elicited cross-reactive Abs. Data are shown for ELISA and neutralization analyses for terminal bleed serum samples (see also Fig. 2). (A) Immunization regimen. Left: Mice were primed at day 0, boosted at day 28, and samples were collected from a terminal bleed at day 77 or 78. Right: Colors used to identify immunizations and symbols indicating a matched (filled in square data points; gray shading around name) or mismatched (unfilled square data points; black outline around name) sarbecovirus antigen. Sarbecovirus strain names are colored in panel B according to clade. (B) RBD-binding ELISA (left) and pseudovirus neutralization (right) results for serum samples from day 77 or 78 after the prime immunization. Immunogens are shown compared with the cohorts immunized with 0.5 µg of an mRNA-based immunogen. Dashed horizontal lines indicate the limits of detection for each assay. Left: Geomeans of ED_50_ values for animals in each cohort (symbols with geometric standard deviations indicated by error bars) are connected by colored lines. Mean titers against RBDs from indicated sarbecoviruses were compared pairwise across immunization cohorts by Tukey’s multiple comparison test with the Geisser-Greenhouse correction (as calculated by GraphPad Prism). Right: Neutralization potencies for serum samples from day 77 or 78 after immunization presented as half-maximal inhibitory dilutions (ID_50_ values) of sera against pseudoviruses from the indicated coronavirus strains. Significant differences between the two cohorts are indicated by asterisks: p<0.05 = *, p<0.01 = **, p<0.001 = ***, p<0.0001 = ****.

**Figure S3.**
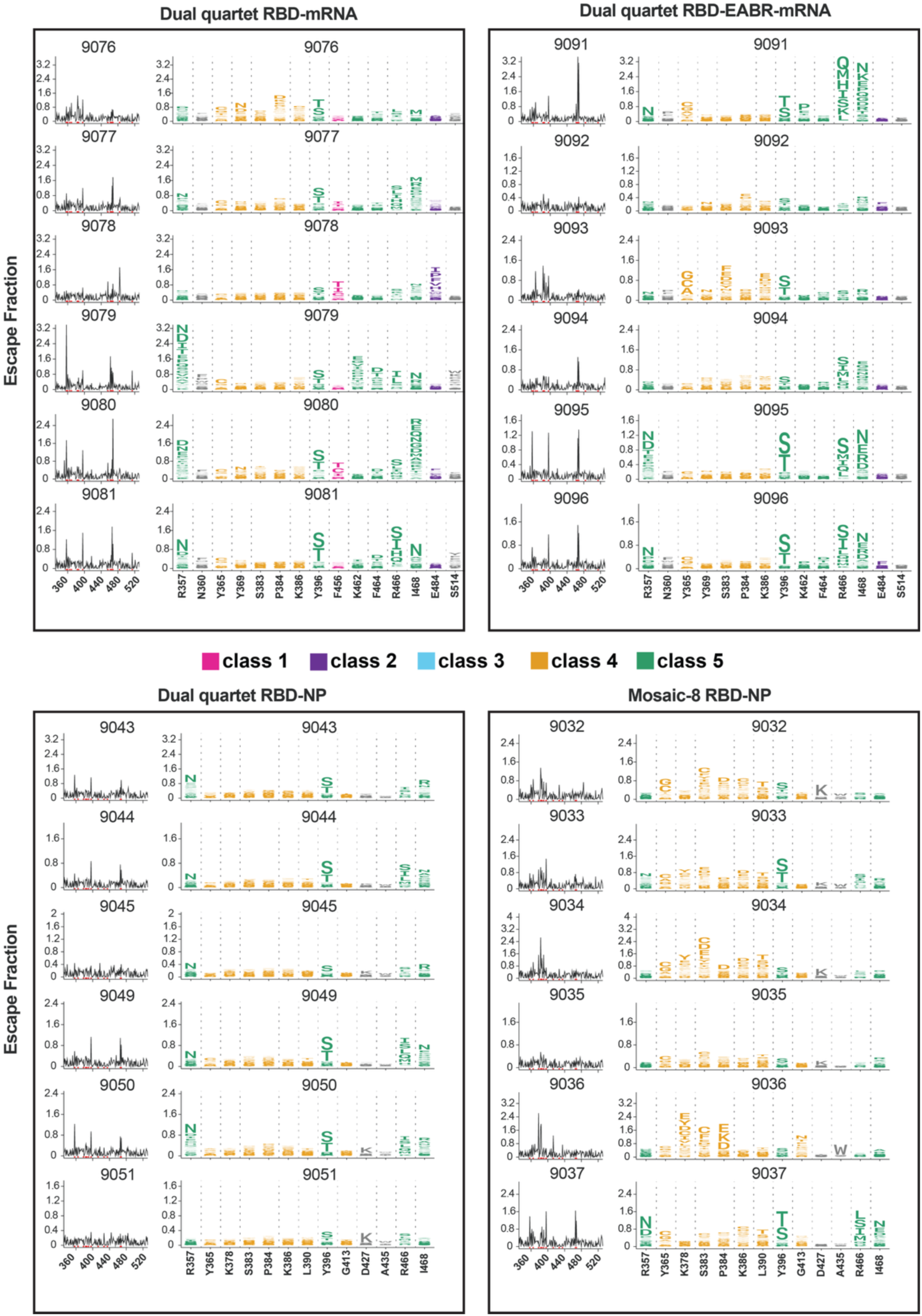
DMS line and logo plots (WA1 RBD library) for individual mice.

**Figure S4.**
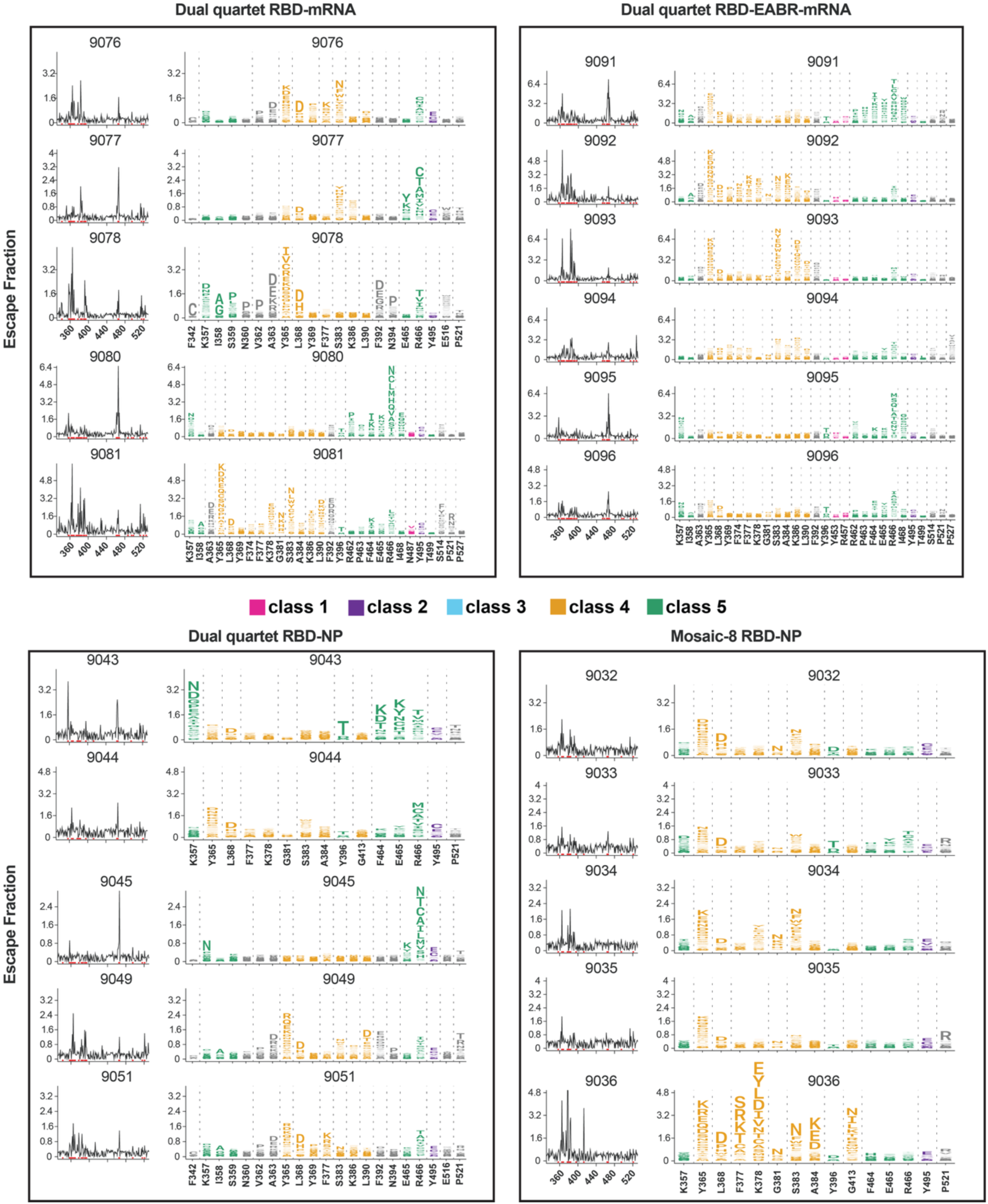
DMS line and logo plots (SARS-1 RBD library) for individual mice.

**Figure S5.**
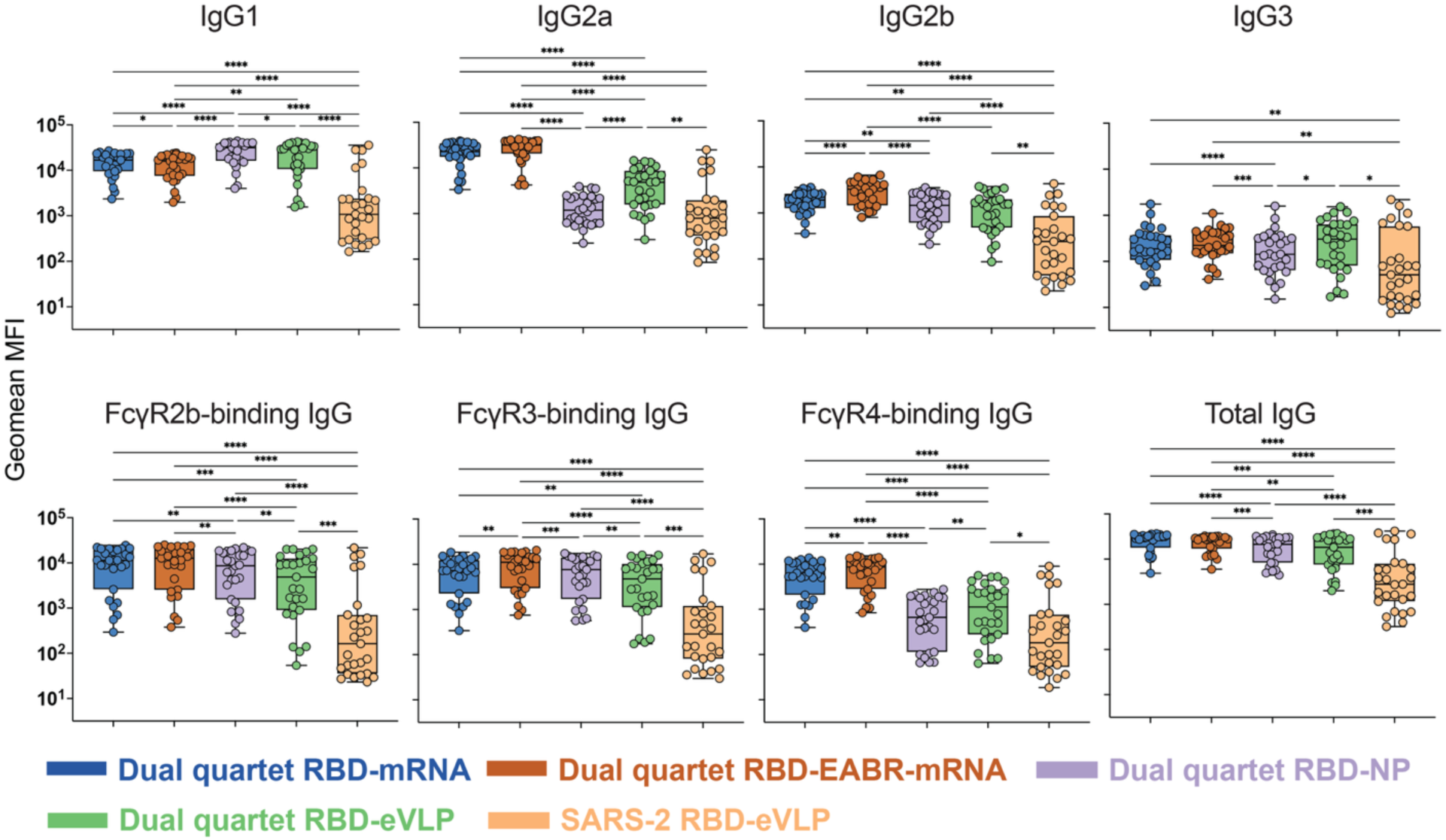
mRNA-encoded dual quartet RBD immunogens elicit balanced IgG subclass and potent FcγR-binding responses. MFI = Median fluorescent intensity. For IgG1, IgG2a, IgG2b, IgG3, FcγR2b-binding IgGs, FcγR3-binding IgGs, FcγR4-binding IgGs, and total IgG, geomean MFI values of the individual responses shown in panel A for each cohort binding to different spikes or RBDs are represented as points in a box and whisker plot and compared pairwise across immunization cohorts by Tukey’s multiple comparison test calculated by GraphPad Prism. Significant differences between cohorts linked by vertical lines in panels B and C are indicated by asterisks: p<0.05 = *, p<0.01 = **, p<0.001 = ***, p<0.0001 = ****. See also Fig. 5.

**Figure S6.**
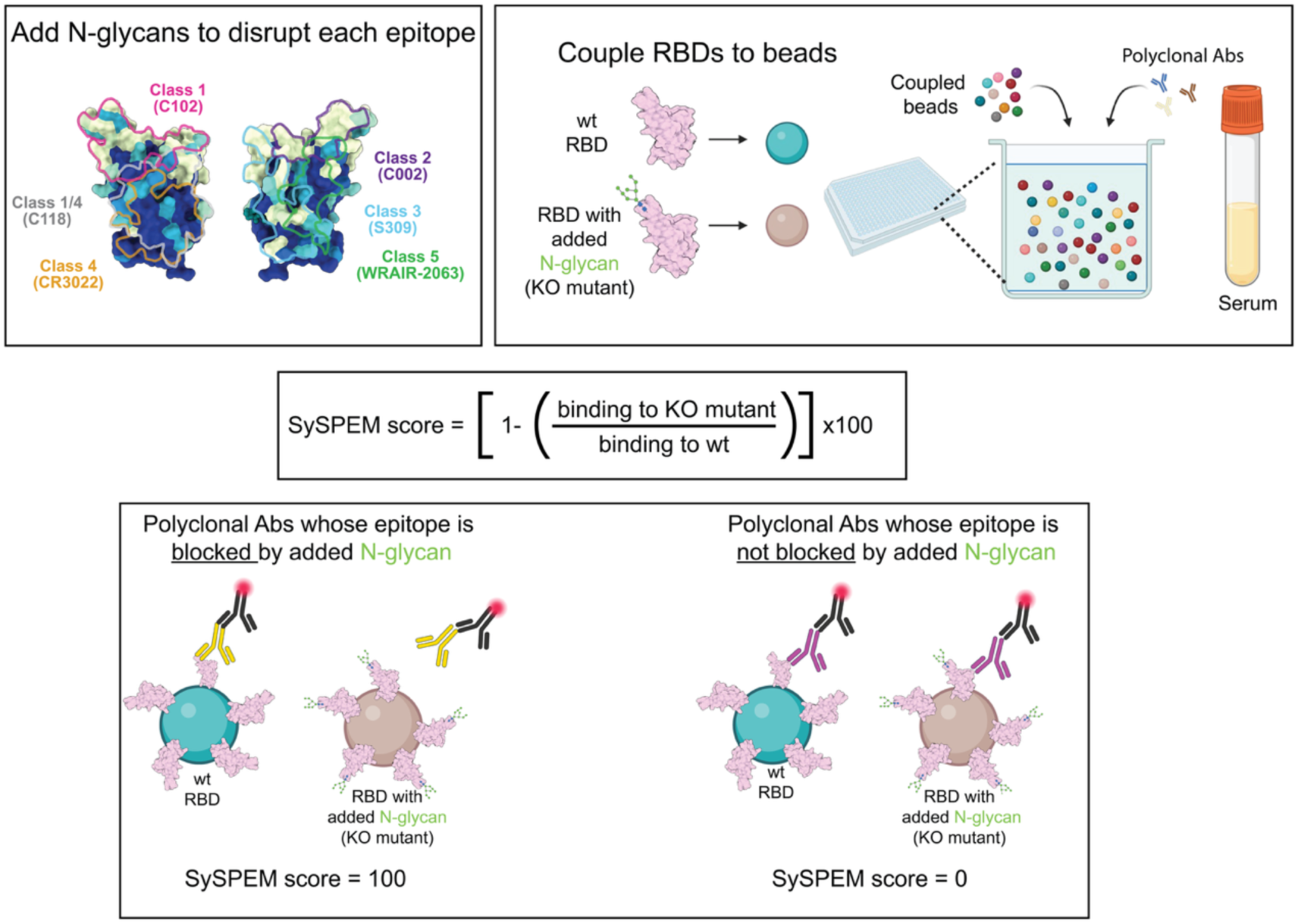
Schematic of SySPEM approach.

**Figure S7.**
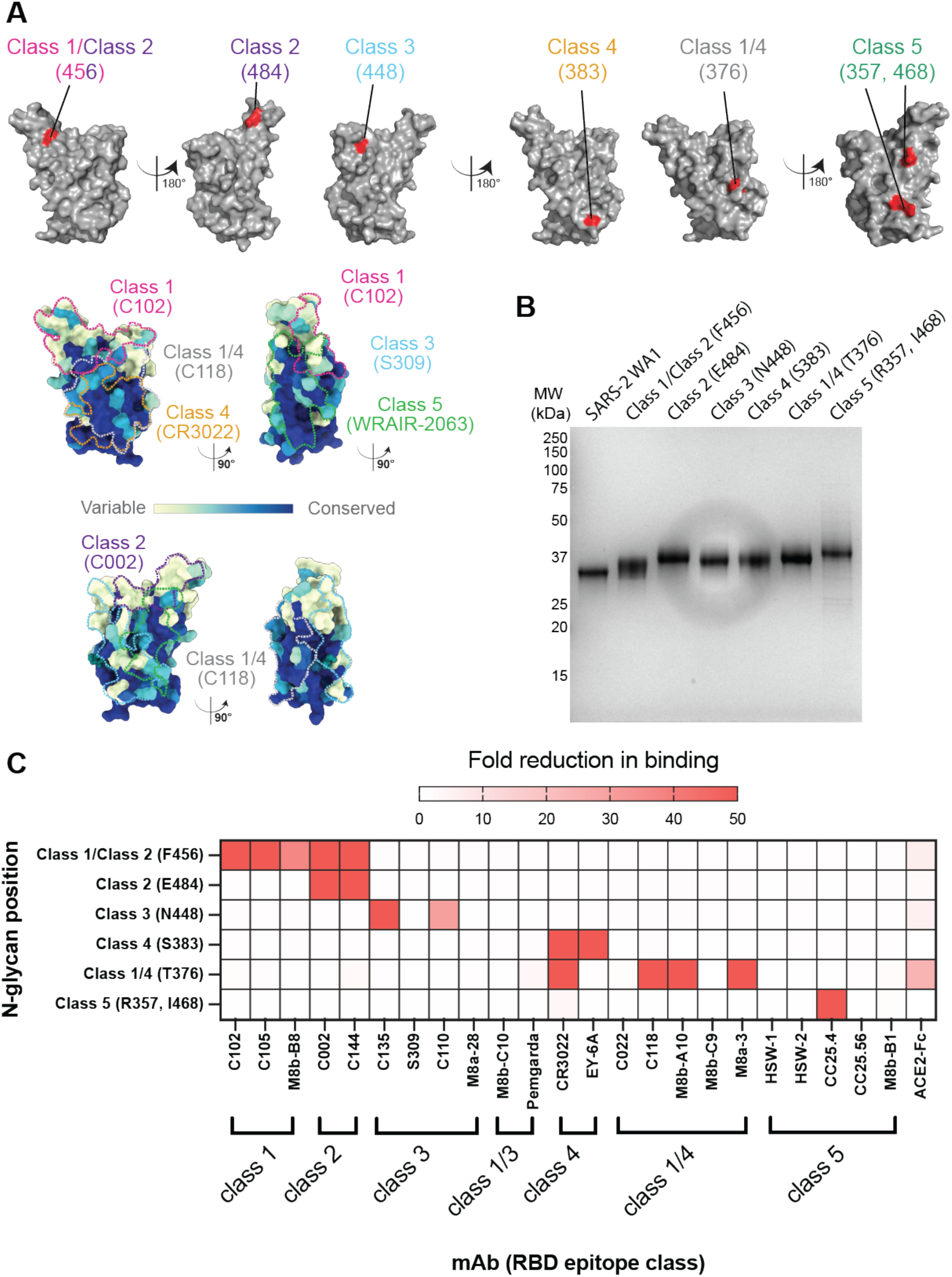
Characterization of RBD N-glycan mutants used for SySPEM analysis. (A) Top: Locations of the added N-glycan(s) in RBD KO mutants, as shown by highlighting of one or more residues that were changed to Asn in the introduced PNGS(s). Bottom: Sequence conservation of 16 sarbecovirus RBDs calculated using ConSurf (Landau et al., 2005) shown on a surface representation of SARS-2 RBD (PDB 7BZ5). Class 1, 2, 3, 4, 1/4, and 5 anti-RBD Ab epitopes (Barnes et al., 2020; Cui et al., 2024; Jensen et al., 2023; Jette et al., 2021) are outlined in dots in different colors using information from representative structures of Abs bound to SARS-2 spike or RBD (C102: PDB 7K8M; C002: PDB 7K8T, S309: PDB 7JX3; CR3022: PDB 7LOP; C118: PDB 7RKV; WRAIR-2063: PDB 8EOO). (B) SDS-PAGE analysis of purified wt RBD and RBD KO mutants. Molecular weight marker positions are shown on the left with the molecular weight indicated in kilodaltons. KO mutants are listed with the RBD epitope class affected by the N-glycan addition(s) and residue number(s) of Asn residue(s) to which N-linked glycan(s) were added. (C) Results of ELISA showing ratio of binding of characterized mAbs or human ACE2-Fc (Jette et al., 2021) to wt RBD versus the RBD KO mutants listed on the left. ELISA EC_50_ values for binding of each reagent to wt RBD and the six RBD KO mutants were derived using, and the fold reduction in binding to each RBD KO mutant was calculated as EC_50_ RBD KO / EC_50_ RBD wt. Classifications of RBD epitopes recognized by the characterized mAbs and ACE-2 Fc are taken from (Barnes et al., 2020; Cui et al., 2024; Fan et al., 2022; Fan et al., 2025b; Jensen et al., 2023; Jette et al., 2021).

**Figure S8.**
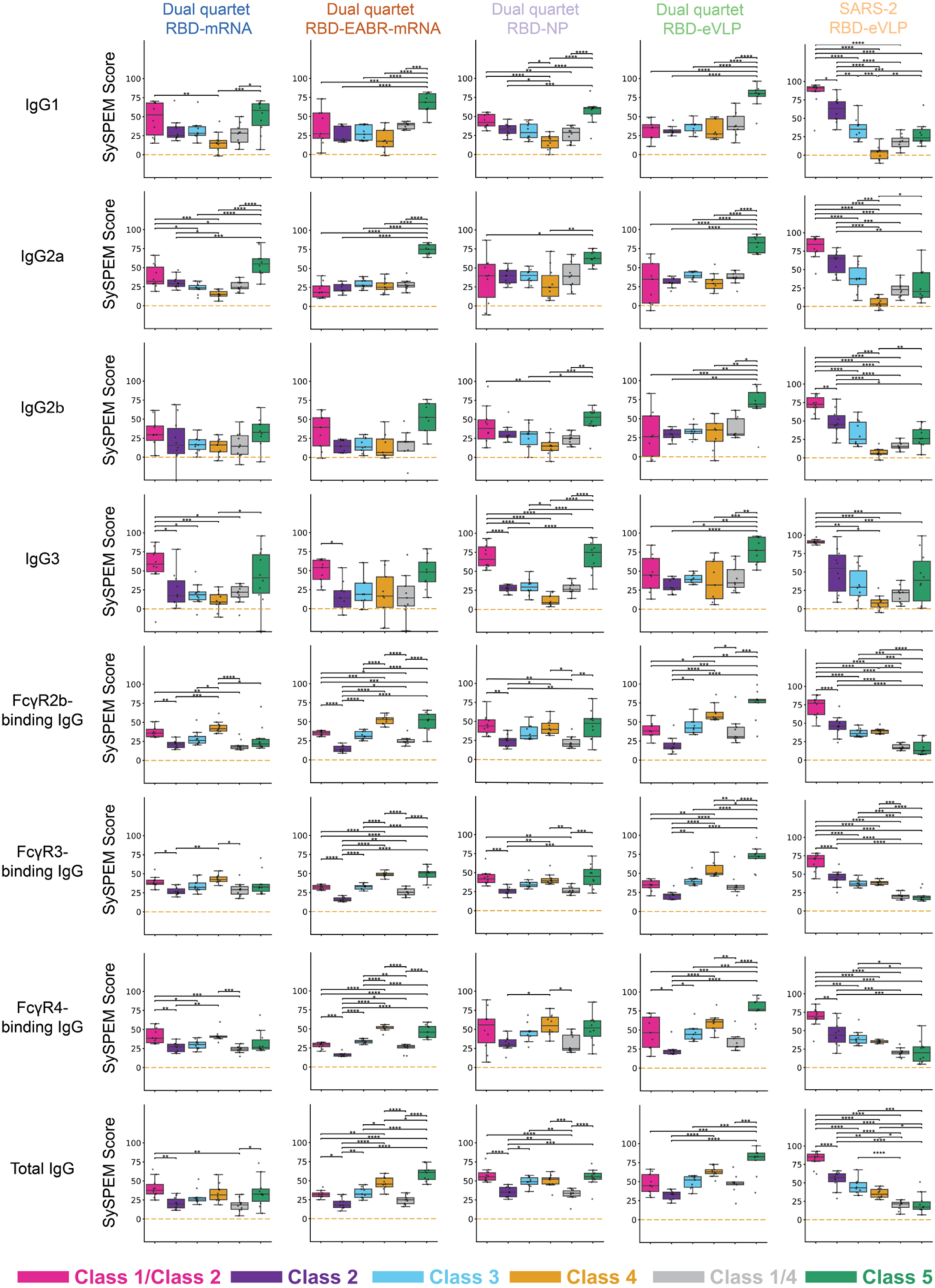
SySPEM score comparisons show distinct epitope profiles across immunogen cohorts. SySPEM scores from individual mice in each immunogen cohort (columns) were determined for each IgG class (rows, IgG subclass plus different FcγR-binding IgGs) with statistical comparisons between targeted epitopes (colors). A SySPEM value of 0 indicates that none of the IgGs in that sample were affected by the glycan addition and therefore the sample did not contain IgGs that recognize that epitope, and a SySPEM value of 100 indicates that all IgGs in that sample recognized that epitope (Fig. S6). A SySPEM value between 0 and 100 indicates the proportion of IgGs in a sample that recognized the epitope that was blocked by glycan addition in the RBD KO mutant. Box and whisker plots of SySPEM scores with individual data points representing one mouse are shown. Significant differences between cohorts were calculated using Tukey’s HSD posthoc test and linked by vertical lines indicated by asterisks: p<0.05 = *, p<0.01 = **, p<0.001 = ***, p<0.0001 = ****. See also Figs. 6 and 7.

## Notes

### Summary of Updates

This version includes some revisions to the abstract and adds some additional discussion which helps to describe the analysis and add additional limitations.

